# The complexities of inferring symbiont function: *Paraburkholderia* symbiont dynamics in social amoeba populations and their impacts on the amoeba microbiota

**DOI:** 10.1101/2021.08.21.457203

**Authors:** James G. DuBose, Michael S. Robeson, Mackenzie Hoogshagen, Hunter Olsen, Tamara S. Haselkorn

**Affiliations:** Department of Biology, University of Central Arkansas, 201 Donaghey Avenue, Conway, AR 72035; Department of Biomedical Informatics, University of Arkansas for Medical Sciences, Little Rock, AR, 72205

**Keywords:** Symbiosis, farming, *Paraburkholderia*, *Dictyostelium discoideum*, amoeba, microbiota

## Abstract

The relationship between the social amoeba *Dictyostelium discoideum* and its endosymbiotic bacteria *Paraburkholderia* provides a model system for studying the development of symbiotic relationships. Laboratory experiments have shown that any of three species of *Paraburkholderia* symbiont allow *D. discoideum* food bacteria to persist through the amoeba lifecycle and survive in amoeba spores, rather than being fully digested. This phenomenon is termed “farming”, as it potentially allows spores dispersed to food poor locations to grow their own. The occurrence and impact of farming in natural populations, however, has been a challenge to measure. Here, we surveyed natural *D. discoideum* populations and found that only one of the three symbiont species, *P. agricolaris*, remained prevalent. We then explored the effect of *Paraburkholderia* on the amoeba microbiota, expecting that by facilitating bacterial food carriage it would diversify the microbiota. Contrary to our expectations, *Paraburkholderia* tended to infectiously dominate the *D. discoideum* microbiota, in some cases decreasing diversity. Similarly, we found little evidence for *Paraburkholderia* facilitating the carriage of particular food bacteria. These findings highlight the complexities of inferring symbiont function in nature and suggest the possibility that *Paraburkholderia* could be playing multiple roles for its host.

## Introduction

Bacterial symbionts have been found to associate with hosts across the tree of life, with often dramatic effects on their ecology and evolution. The roles of bacterial symbionts within their hosts can range from mutualism to parasitism and can vary based on host and environmental contexts, as well as on the length of the association and mode of transmission (R. M. Anderson & May, 1982; Ewald, 1987; Moran, 2006). Amoebae, a polyphyletic group of unicellular eukaryotes, serve as promising model systems for studying symbiotic relationships, particularly the bacterial transition from free-living to intracellular. When amoebae feed on bacteria, a selective pressure is exerted that favors bacteria with mechanisms to resist digestion (Brüssow, 2007; Dunn et al., 2018; Sun et al., 2018). Hence, amoebae serve as intracellular training grounds for bacteria to evolve phagocytosis resisting mechanisms, allowing certain bacteria to occupy amoebae as endosymbionts (Casadevall, 2008; Greub & Raoult, 2004; Molmeret et al., 2005). Bacterial endosymbionts have been commonly identified in *Acanthamoeba* and *Vermamoeba* hosts, including lineages of Chlamydiae, *Proteobacteria, Bacteroidetes*, and *Dependentiae* (Delafont et al., 2015; Horn & Wagner, 2004; Schmitz-Esser et al., 2008), although the effects on their hosts are often unclear. For example, while some Chlamydiae symbionts live as energy parasites that are costly to their hosts (Horn, 2008), other Chlamydiae species provide benefits by protecting *Acanthamoeba* from the pathogenic *Legionella* bacteria and enhancing cell motility and growth. (Ishida et al., 2014; König et al., 2019; Maita et al., 2018; Okude et al., 2012). The difficulty of studying the microbial community dynamics in natural environments dampens our understanding of the ecological contexts of amoeba – bacteria interactions, which makes inferring the function of a particular symbiont in its host a challenge.

The social amoeba *Dictyostelium discoideum* has gained traction as a model organism to study host cell – bacteria interactions in laboratory settings. Social amoebae are soil-dwelling protists that have unique life cycles that involve the transition from a single-cellular state to a multicellular state when their bacterial food source is depleted (Bozzaro, 2019). Tens of thousands of amoeba cells aggregate to ultimately form a fruiting body, which consists of two primary structures: the sorus, a structure used to house spores, and the stalk, which is composed of dead cells that differentiated to support the sorus (Kessin, 2001). It was previously thought that sentinel cells, which perform immune-like functions, cleared all bacteria from the amoeba fruiting body throughout the social cycle (G. Chen et al., 2007), as evidenced by the inability to culture bacteria from many laboratory-isolated *D. discoideum* clones. Recent studies, however, have shown that in nature *D. discoideum* harbors microbiota consisting of both transient and persistent bacterial symbionts (Brock et al., 2018; Sallinger et al., 2021), belonging to the phyla Proteobacteria, Actinobacteria, Bacteroidetes, Firmicutes, Chlamydiae, and Acidiobacteria. This recent characterization of the *D. discoideum* microbiota has expanded its potential to serve as a model system for studies pertaining to microbiome formation and maintenance (Farinholt et al., 2019) and some evidence suggests that *D. discoideum* may play an active role in its microbiota formation by secreting lectin proteins that bind bacteria during the social cycle (Dinh et al., 2018).

In natural populations of *D. discoideum*, the most widespread and persistent bacterial symbionts are *Paraburkholderia*, Chlamydiae, and *Amoebophilus* (Haselkorn et al., 2021). Although the functions of these symbionts in natural populations are largely unknown, the *Paraburkholderia* farming symbionts have been the best characterized. The phenomenon of farming was demonstrated primarily in laboratory experiments by comparing the ability of *Paraburkolderia* infected and uninfected *D. discoideum* to grow when a sorus was plated without food bacteria. *Paraburkholderia* are inedible, but they allow bacterial food carriage that is presumed to benefit the amoeba in nature if the spores are dispersed to an area that is scarce in food bacteria, as the soil could be seeded with the carried bacteria (Brock et al., 2011; DiSalvo et al., 2015; Haselkorn et al., 2019). There is, however, a cost to infection, as *Parburkholderia*-infected *D. discoideum* produce fewer spores in food rich laboratory conditions. While many members of the *Paraburkholderia* and closely related *Burkholderia* genera form symbiotic interactions with a variety of plant, fungi, and insect hosts (Esmaeel et al., 2018; Kaltenpoth & Flórez, 2020; Nazir et al., 2014), only three *Paraburkholderia* species, *P. agricolaris, P. bonniea*, and *P. hayleyella* have been found to confer farming and form persistent symbiotic relationships with *D. discoideum* (Haselkorn et al., 2019). The variability in fitness costs and the extent of food carriage that each symbiont permits makes this *Paraburkholderia* – *D. discoideum* symbiosis context dependent (Khojandi et al., 2019), like many symbiotic relationships (Oliver et al., 2008). Screenings for *Paraburkholderia* in natural *D. discoideum* populations have shown that *P. agricolaris* tends to be the most prevalent, followed by *P. hayleyella* and *P. bonniea* (Haselkorn et al., 2019). Indeed, these differences in symbiotic potential likely translate to different host outcomes, which could be responsible for the differing levels of prevalence between each *Paraburkholderia* species.

Measurements of fitness in laboratory settings have aided our understanding of the consequences of *Paraburkholderia* infection, but its role in natural populations is less clear. To better understand the occurrence and impact of this of this symbiosis, we used PCR based screenings of *D. discoideum* populations to identify current symbiont prevalence, transmission patterns, and genetic diversity. Since symbiont transmission modes play a large role in determining whether a symbiont’s evolutionary trajectory will lead to mutualism, benign parasitism, or extreme pathogenicity (R. M. Anderson & May, 1982), we looked for evidence showing whether *P. agricolaris* was environmentally acquired or vertically transmitted between social amoeba hosts. To investigate how *Paraburkholderia* might be conferring bacterial food carriage in a natural setting, we compared the microbiota of infected and uninfected *D. discoideum*. We hypothesized that if *Paraburkholderia* was indiscriminately facilitating the carriage of food bacteria, infected amoeba would possess more diverse microbiota. Alternatively, if *Paraburkholderia* was facilitating the carriage of certain food bacteria, perhaps indicative of *D. discoideum* feeding preferences, we hypothesized that *Paraburkholderia* infected amoeba would have microbiota that were similar in composition to each other. Inconsistent with our hypotheses, we found that *Paraburkholderia* tended to dominate the microbiota, often correlating with a decrease in microbiota diversity. Overall, our results provide insight into the natural history of the *Paraburkholderia* – *D. dicoideum* relationship and further our understanding of the ecological relevance of the farming phenotype.

## Methods

### Soil Collection and Amoeba Culturing

We re-sampled natural *D. discoideum* populations in the Virginia Mountain Lake area (Haselkorn et al., 2019, 2021). Soil, sampled from below the leaf litter layer of the forest floor or below rotting logs, was collected from ten different sites at each of four separate locations (Supplemental Table 1) and was plated on hay agar within 96 hours (Supplemental Table 2). After 4-5 days of growth time, Dictyostelid species were morphologically identified via stereomicroscopy (Raper, 2014), and 5-10 sori from each patch were collected with sterile pipette tips and suspended into 250µL of KK2 spore buffer. Every patch of *D. discoideum*, identified by the presence of a basal disc at the base of its stalk, was collected, as well as other social amoeba species, including *D. mucoroides, D. giganteum, D. purpureum, Heterostelium pallidum*, and *Polysphondylium violaceum*. We then resuspended fruiting bodies in 100µL of KK2 buffer for “passaging”: replating onto SM/5 agar (Supplemental Table 2) with the addition of 200µL of a 1.5 OD_600nm_ suspension of *Klebsiella pneumoniae* food bacteria to increase the quantity of fruiting bodies. Subsets from each amoeba species were sequenced to confirm amoeba identity (See following section).

### Symbiont Screening

DNA from 15-20 fruiting bodies from the passaged SM/5 plates was extracted using the a Chelex/proteinaseK protocol as in (Haselkorn et al., 2019). PCRs were conducted to screen each sample for *Dictyostelium* DNA by amplifying approximately 500 base pairs of the 18S rRNA gene (Baldauf et al., 2018). To screen for *Paraburkholderia*, we used the *Paraburkholderia* specific primers that amplify a small section of the 16S rRNA gene (Salles et al., 2002). PCR primers and amplification conditions are listed in Supplemental Table 3. Each PCR screen included a positive control, two negative PCR controls, and two negative DNA extraction controls to ensure there was no contamination. Positive samples were sent for Sanger sequencing at Eurofins Genomics. Sequences were trimmed using Geneious v8 (http://www.geneious.com) (Kearse et al., 2012), and NCBI Blast was used to classify *Paraburkholderia* species. In samples that were identified as either *P. agricolaris, P. bonniea*, or *P. hayleyella*, the LepA gene, which allows for better species level identification and haplotype resolution (Haselkorn et al., 2019), was then amplified and sequenced (Spilker et al., 2009). Additionally, we screened for Chlamydiae and *Amoebophilus* using symbiont specific primers for each (Ossewaarde & Meijer, 1999) (Supplemental Table 3).

### *P. agricolaris* phylogeny, Population Genetics, and Co-infection Statistics

LepA sequences were used to construct a phylogenetic tree of *P. agricolaris* haplotypes. Overlapping peaks in the DNA chromatogram at a single site were indicative of co-infections with multiple haplotypes and were excluded from analysis because haplotype designations were not able to be resolved. A Maximum Likelihood phylogenetic tree was constructed in MEGA7 using the Tamura 3-parameter+G+I model with 1 000 bootstrap replicates (Kumar et al., 2016). Previously identified LepA haplotypes were also placed to determine if these haplotypes had previously been identified (Haselkorn et al., 2019).

To examine *P. agricolaris* genetic differentiation between *D. discoideum* populations, pairwise F_ST_ values were calculated in DNAsp (v6) (Rozas et al., 2017) using non-coinfected LepA sequences (n=65). Concatenated 16S rRNA and LepA sequences were then used to look for evidence of recombination using the Recombination Detection Program (RDP) (v4) (Martin et al., 2015). Pairwise Fisher exact tests (2×2 contingency tables) between *P. agricolaris*, Chlamydiae, and *Amoebophilus*, were used to investigate ecological co-infection tendencies between symbionts.

### Laboratory inoculations of *D. discoideum* with different *Paraburkholderia* species

Separate *P. agricolaris, P. bonniea*, and *P. hayleyella* solutions were made by resuspending stationary phase bacterial colonies from SM/5 plates into KK2 buffer at a concentration of 1.5 OD_600nm_ and combined with 1.5 OD_600nm_ *K. pneumoniae* suspensions at a final volume of 95%/5% *K*.*pneumoniae*/*Paraburkholderia*. We then plated 10^5^ *D. discoideum* spores (strain QS864) with 200µL of each *Paraburkholderia* solution onto SM/5 plates and incubated at room temperature. Amoebae were collected after one week of growth time, and *Paraburkholderia* was cultured from fruiting bodies to confirm infection. Ten fruiting bodies from each plate were resuspended in KK2 spore buffer and passaged again to confirm persistent *Paraburkholderia* infection. *Paraburkholderia* infected *D. discoideum* lines were then inoculated into 2.5 grams of local soil collected from a natural area maintained by the Department of Biology at the University of Central Arkansas (Conway, AR). Soil contents were subsequently plated as describe above, and *D. discoideum* fruiting bodies were collected directly from these soil pates after one week of growth time.

### DNA Extractions for Microbiota Analyses

Approximately 50-100 sori were collected from passaged SM/5 plates (natural infections) or soil plates (lab infections), placed into Hyclone water, and stored at −80°C until DNA was extracted. DNA extractions were performed using the Qiagen DNeasy PowerSoil Pro Kit and the Qiagen DNeasy UltraClean Pro Kit, according to manufacturer’s protocols. Samples were then sent for sequencing to Argonne National Laboratories (Lemont, IL, USA). Here, samples were sequenced on the Illumina MiSeq platform using 250bp paired-end cycles. Library preparation and sequencing was performed using the 799F-1115R primer set, which amplifies the V5-V6 region of the 16S rRNA gene (Redford et al., 2010), as the more commonly used primers for the V4 region preferentially amplify amoeba mitochondrial DNA (Sallinger et al., 2021). Sequences were returned as FASTQ files with adaptors removed.

### Illumina Sequence Processing

Illumina sequence files were demultiplexed and analyzed in QIIME2 (v2020.6). Chimeric sequences were removed, primers were trimmed, and paired-ends were joined using q2-dada2 plugin (Callahan et al., 2016; Johnson et al., 2008; McDonald et al., 2012; Sayers et al., 2021). The q2-RESCRIPt plugin was used to curate the SSU SILVA NR99 (version 138) database, and to train a custom V5-V6 amplicon specific classifier (Robeson et al., 2020). The classify-sklearn method was used in the q2-feature-classifier plugin to assign taxonomic names to amplicon sequence variants (ASVs) (Bokulich et al., 2018; Janssen et al., 2018; McKinney, 2010). ASVs classified as either mitochondria or chloroplast, as well as ASVs that did not classify to the phylum-level were removed for downstream analysis. ASVs with a percent identity/alignment of less than 0.80/0.80 to the SILVA reference sequences were excluded using the q2-quality-control plugin (Camacho et al., 2009). A phylogenetic tree for downstream analyses was created by inserting ASVs into the Silva 128 reference database using SATé-enabled phylogenetic placement (SEPP), where all fragments were successfully inserted (Janssen et al., 2018). Although the V5-V6 region of 16S rRNA did not provide species level resolution, we identified the genus *Burkholderia-Caballeronia-Paraburkholderia* to represent *P. agricolaris* based on Sanger sequences of the more variable LepA gene in our naturally infected study. In laboratory inoculated samples, the genus *Burkholderia-Caballeronia-Paraburkholderia* was identified to represent both *P. agricolaris* and *P. hayleyella* and the genus *Pandorea* was identified to represent *P. bonniea*, based on their high abundance in their respective infection groups.

### Microbiota Diversity Analyses

After constructing alpha-rarefaction curves, natural samples were rarefied to a depth of 3 950 and laboratory inoculated samples were rarefied to a depth of 4 000 (Weiss et al., 2017). This allowed us to retain 28 *P. agricolaris* infected 16 uninfected natural samples and 19 *P. agricolaris*, 9 *P. bonniea*, and 10 *P. hayleyella* artificially infected samples. Alpha-diversity was evaluated by using the q2-diversity plugin to calculate average ASVs, Pielou’s evenness, and Faith’s phylogenetic diversity (Janssen et al., 2018; Pielou, 1966). Significance of each metric was evaluated using Kruskal-Wallis and pairwise Kruskal-Wallis tests (α=0.05 with FDR q-values) (Kruskal & Wallis, 1952). Beta-diversity was evaluated by using the q2-deicode plugin to calculate the Aitchison distances between non-rarefied sample groups (Martino et al., 2019).

Weighted UniFrac and Unweighted UniFrac were also used on rarefied data to evaluate beta-diversity using the q2-diversity plugin (Chang et al., 2011; J. Chen et al., 2012; Faith et al., 1987; C. A. Lozupone et al., 2007; C. Lozupone & Knight, 2005). Ordination plots were generated for each beta-diversity metrics and between group significance was evaluated using PERMANOVA and PERMDISP with 999 permutations and FDR adjusted p-values (M. J. Anderson, 2001). When PERMAONVA and PERMDISP results were both significant, ordination plots were consulted to determine separation between groups. The feature table generated by QIIME2 was uploaded to microbiomeanalyst.ca to perform co-association analyses using the SparCC method.

## Results

### *P. agricolaris* remains prevalent in natural *D. discoideum* populations but does not co-associate with Chlamydiae or *Amoebophilus* endosymbionts

To identify the current prevalence of *D. discoideum* endosymbionts, we screened 244 *D. discoideum* isolates collected from the Mountain Lake area in 2019 for *Paraburkholderia*, Chlamydiae, and *Amoebophilus*. We then compared the prevalence of each endosymbiont in 2019 to that of a previous collection in 2000 (Haselkorn et al., 2019, 2021). *P. agricolaris* exhibited a prevalence of 35.1%, up from a 14.9% prevalence in 2000. *P. bonniea* was only found once in our sampling, resulting in a prevalence of 0.79% (down from 2.6% in the 2000 collection). *P. hayleyella*, which was previously identified to be 8% prevalent in 2000, was not found in our collection. *P. agricolaris* appeared to be stochastically distributed between each location and collection site, where prevalence varied from less than 15% at location 1 to greater than 60% at location 4 (Figure 1, Supplemental Table 2). Chlamydiae, which was 19% prevalent in 2000, was found to be 58% prevalent in our collection. *Amoebophilus*, 37% prevalent in 2000, was found to be 18% prevalent in our collection. Pairwise Fishers exact tests were used to look for signs of positive or negative co-associations between *P. agricolaris*, Chlamydiae, and *Amoebophilus* at each location. A positive association was observed between *P. agricolaris* and Chlamydiae at location 3 (p=0.0472, 23 expected with 26 observed); however, no other significant associations were found (Supplemental Table 4).

**Figure 1.**
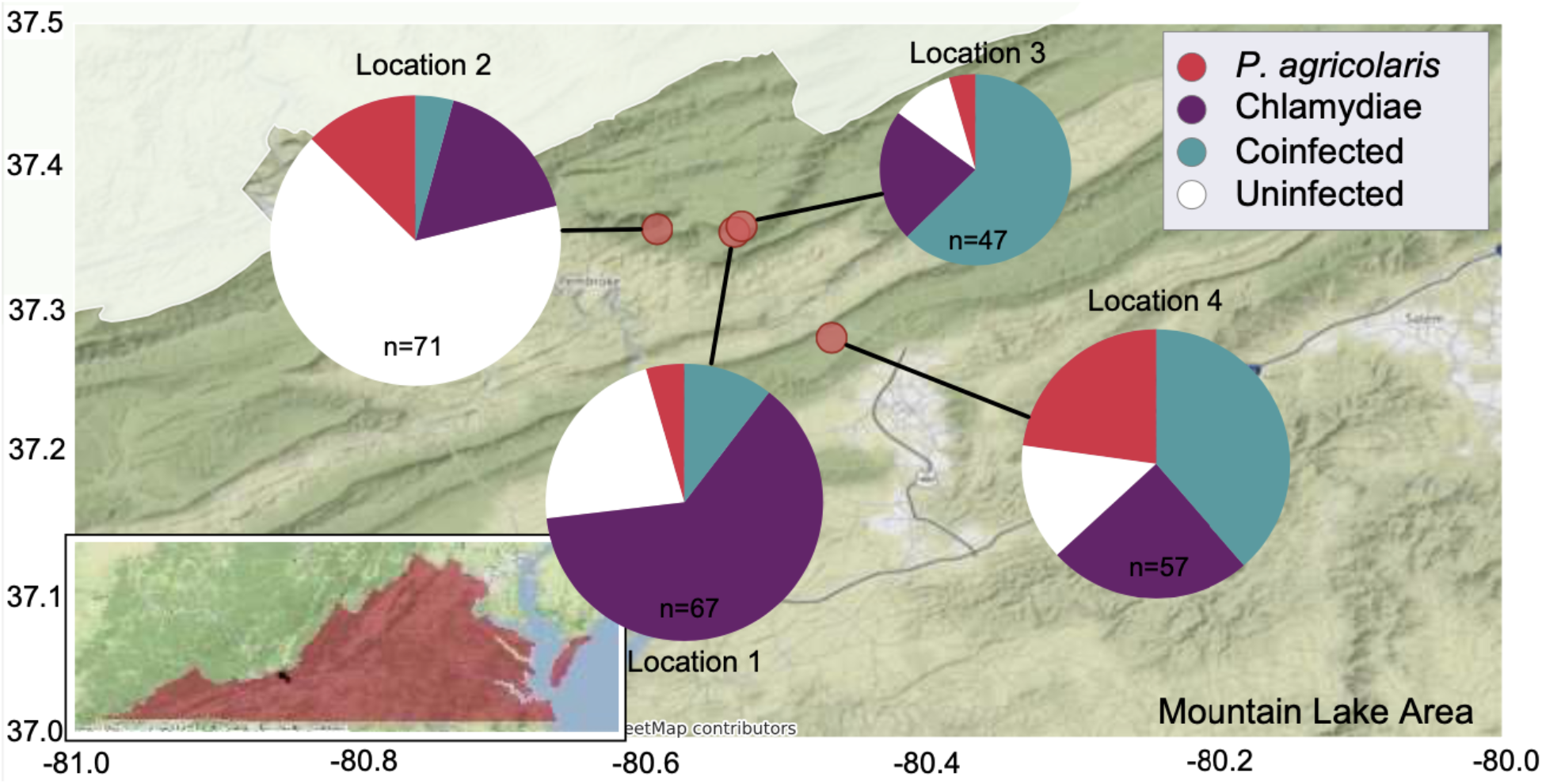
*Paraburkholderia* and Chlamydiae are prevalent in natural *D. discoideum* populations. Pie charts indicate single and co-infection prevalence rates for each site sampled in the Mountain Lake, Virginia area.

### *P. agricolaris* is genetically diverse and shows evidence of recombination and environmental acquisition

To gain insight into the natural and co-evolutionary history of *P. agricolaris* and *D. discoideum*, we investigated *P. agricolaris* haplotype diversity, genetic recombination tendencies, and transmission patterns. Phylogenetic analyses of LepA sequences (n=65) revealed ten distinct haplotypes, and placement of previous *P. agricolaris* sequences showed that 7 of these haplotypes have not been previously identified (Figure 2a) (Haselkorn et al., 2019). Haplotype 7 was the only haplotype found in amoebae from multiple locations and pairwise F_ST_ comparisons showed that the locations harboring haplotype 7 were the most similar in terms of *P. agricolaris* genetic composition (Supplemental table 4). Furthermore, up to 5 different haplotypes were identified in amoebae from a single soil sample (Supplemental Table 5). Using concatenated 16S rRNA and LepA sequences we identified two putative inter-locus recombination events, both between haplotypes 2 and 9. We used the presence of a *P. agricolaris* haplotype infecting multiple amoebae species as evidence of environmental acquisition. In addition to infecting *D. discoideum*, haplotype 3 was identified in *D. purpureum*, haplotype 4 was identified in *D. mucoroides*, haplotype 8 was identified in *H. pallidum*, and haplotype 7 was identified in *D. giganteum, H. pallidum, P. violaceum*, and *D. mucoroides* (Figure 2).

**Figure. 2.**
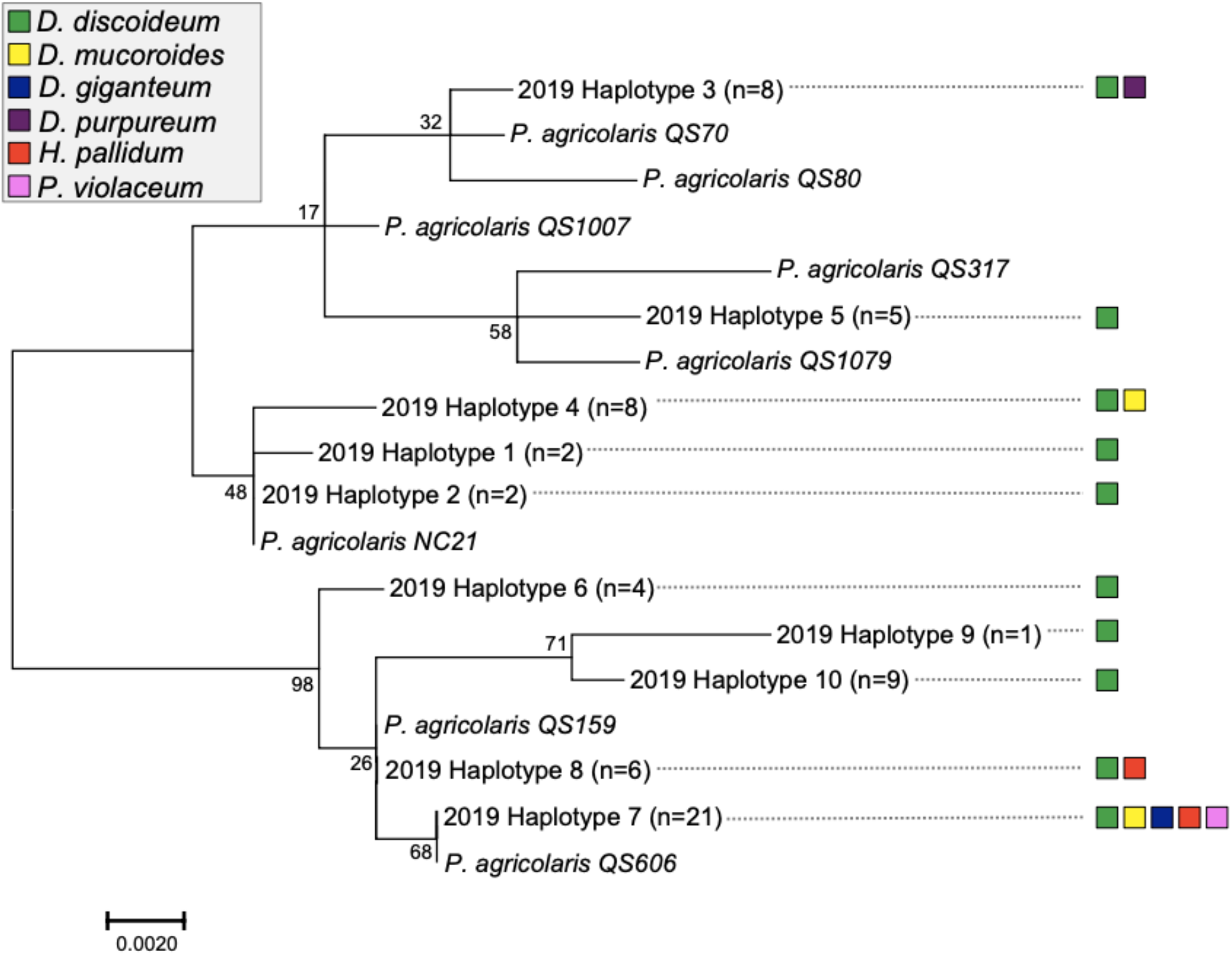
Phylogenetic evidence for environmental acquisition of *P. agricolaris*. Maximum likelihood phylogenetic tree of *P. agricolaris* haplotypes constructed from the LepA housekeeping gene. Colored boxes represent every amoeba species found to contain the corresponding haplotype. Multiple amoebae species harboring the same haplotype was used as indication of environmental acquisition of *P. agricolaris*.

### *Paraburkholderia* symbionts dominate the *D. discoideum* microbiota and infection correlates with a general decrease in microbiota diversity

Our investigation of the impacts of *Paraburkholderia* on the natural *D. discoideum* microbiota was limited to *P. agricolaris*, as *P. bonniea* and *P. hayleyella* were not found. Overall, denoising of 16S rRNA sequences resulted in 130 amplicon sequence variants (ASVs) with a total frequency of 1 573 424 reads. These were represented by 6 phyla, 9 classes, 33 orders, 49 families, and 57 genera. The majority of sequences belonged to the genus *Burkholderia-Caballeronia-Paraburkholderia* (57.16%), which was identified to represent *P. agricolaris* based on Sanger sequences of the more variable LepA gene. Although we did not find *P. hayleyella* or *P. bonniea* infected *D. discoideum* in nature, we were able to gain insights on their impacts by culturing artificially infected *D. discoideum* on local soil. This yielded 168 ASVs with a total frequency of 2 135 808 reads. These were represented by 10 phyla, 15 classes, 49 orders, 77 families, and 140 genera.

If *Paraburkholderia* infection was somehow facilitating the carriage of random environmental bacteria, we posited that *Paraburkholderia*-infected *D. discoideum* would have a more diverse microbiota than uninfected ones. However, we found that *P. agricolaris* tended to dominate the *D. discoideum* microbiota, accounting for an average of 71.35% of bacterial community abundance, for greater than 90% of reads in the majority (16/28) of infected samples, and for 76.86% of all reads (Figure 3a). The *D. discoideum* microbiota evenness and richness was not significantly altered by *P. agricolaris* infection (Pielou’s Evenness, p=0.213399; observed ASVs, p=0.376987). However, *D. discoideum* infected with *P. agricolaris* did possess less phylogenetically diverse microbiota than uninfected *D. discoideum* (Faith’s Phylogenetic Diversity, p=0.048117) (Figure 4a). Laboratory inoculation of each *Paraburkholderia* species into *D. discoideum* yielded similar results. *P. agricolaris, P. bonniea*, and *P. hayleyella* each had abundances of greater than 90% in 13/19, 8/9, and 10/10 infected *D. discoideum* isolates, respectively (Figure 3b). Consistent with the trend in *Paraburkholderia* abundance, laboratory inoculation with each of the *Paraburkholderia* species resulted in significant reductions in *D. discoideum* microbiota evenness, richness, and phylogenetic diversity (Pielou’s Evenness, p ≤ 0.001515; observed features (ASVs), p ≤ 0.041061; Faith’s Phylogenetic Diversity, p ≤ 0.041759) (Figure 4b). *P. hayleyella* infection significantly reduced microbiota evenness when compared to *P. agricolaris* (p = 0.041457), but no other pairwise differences were observed between each *Paraburkholderia* species (Figure 4b).

**Figure. 3.**
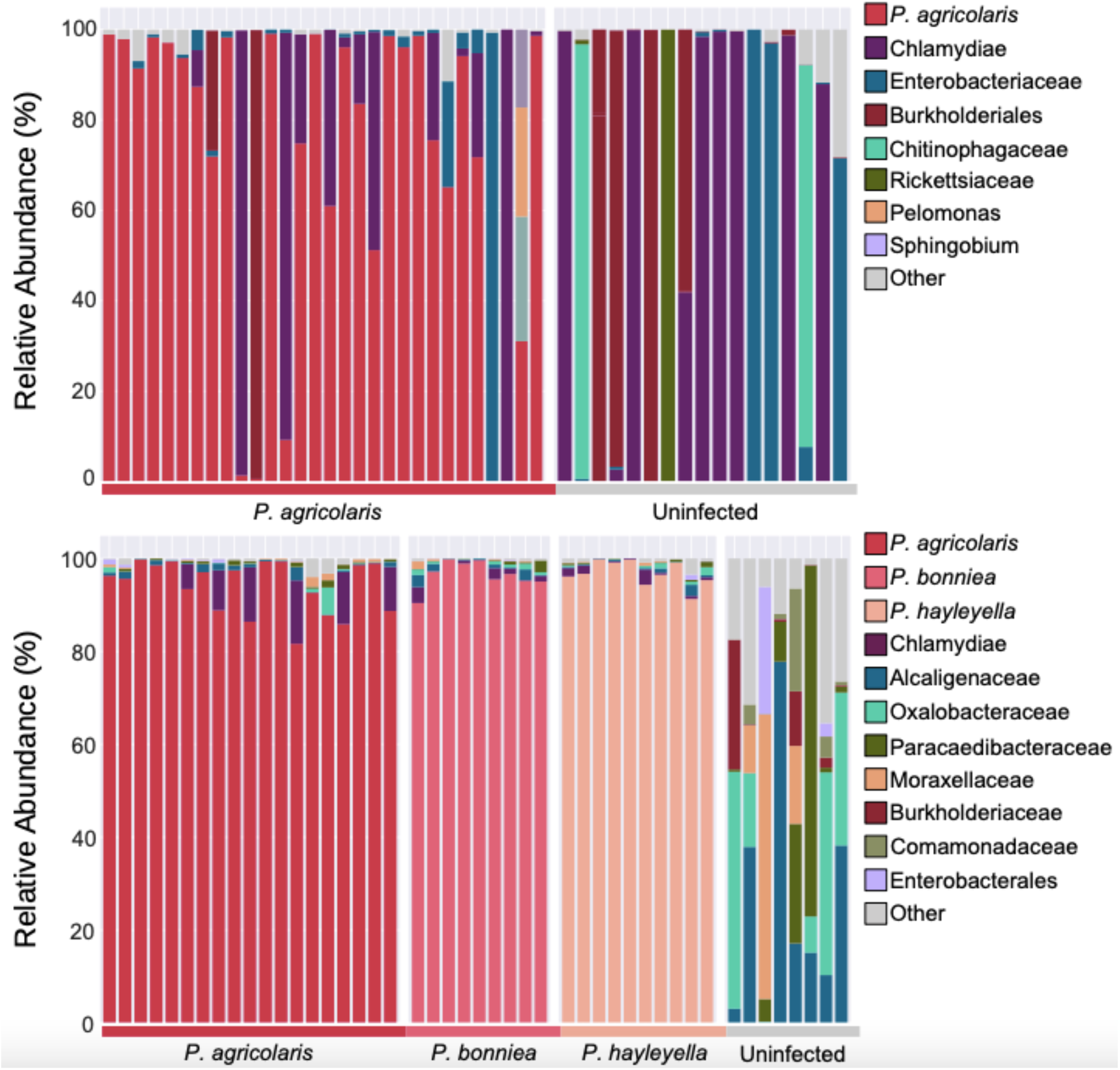
*Paraburkholderia* infectiously dominates the *D. discoideum* microbiota. Bar plots show the members of the most abundant taxa within the microbiota of A) Naturally infected *D. discoideum* isolates and B) artificially infected *D. discoideum* isolates.

**Figure. 4.**
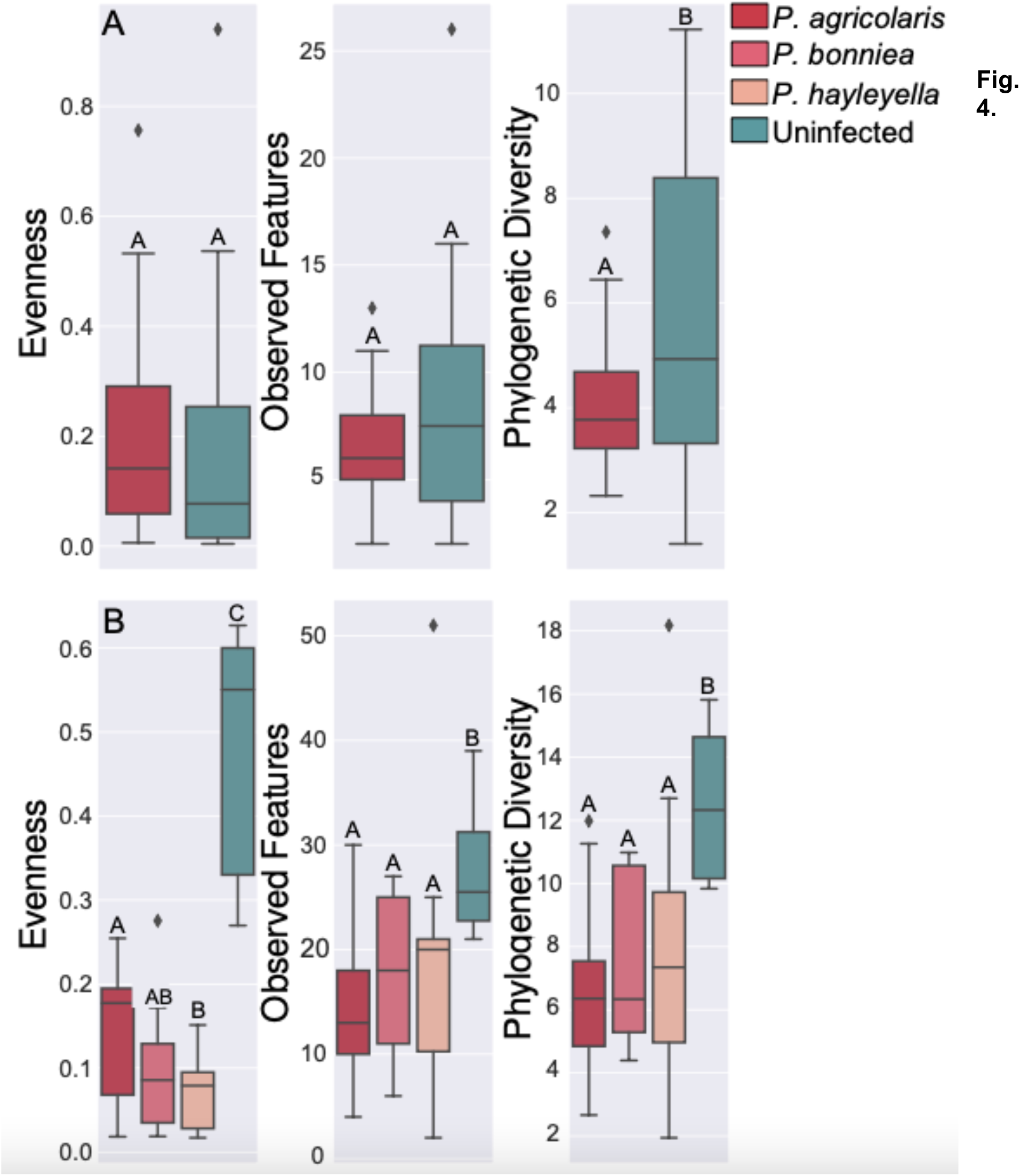
*Paraburkholderia* does not diversify the *D. dicoideum* microbiota. Alpha-diversity box plots showing the Pielou’s evenness, observed ASVs, and Faith’s phylogenetic diversity for A) isolates naturally infected with *Paraburkholderia* and B) isolates artificially infected with each *Paraburkholderia* symbiont. Boxes span the first and third quartiles and whiskers represent ± 1.5 IRQ. Letters shared in common denote no significant difference (Kruskal-Wallis α=0.05 after FDR correction.

### Overabundance of *Paraburkholderia* alters the composition and structure of the *D. discoideum* microbiota

While *Paraburkholderia* infection did not diversify the *D. discoideum* microbiota, we thought it possible that *Paraburkholderia* might facilitate the carriage of specific food bacteria, altering the microbiota structure and/or composition. ß-diversity comparisons using weighted UniFrac, which measures microbial community structure by considering the phylogenetic relatedness and relative abundances of each ASV, showed that the microbiota of infected *D. discoideum* were similar to each other, and dissimilar to the microbiota of uninfected *D. discoideum* (PERMANOVA, pseudo-F=10.5405, p=0.001; PERMDISP, F=8.97892, p=0.01; where significant PERMDISP results were negated by the clear grouping of *P. agricolaris* infected microbiota in PCoA space) (Figure 5a). This is not surprising given the high relative abundance of *Parburkholderia*. ß-diversity comparisons using unweighted UniFrac, which measures microbial community composition by only considering phylogenetic relatedness and presence or absence of ASVs showed the effect of *Paraburkholderia* without directly considering its relative abundance. Here we also saw significant differences between *P. agricolaris* and infected and uninfected amoebae (PERMANOVA, pseudo-F=3.3698, p=0.002; PERMDISP, F=14.0089, P=0.003 where significant PERMDISP results were negated by a clear distinction between infected and uninfected samples upon PCoA visualization). (Figure 5b). The impact, however, is unlikely due to carriage of specific food bacteria, but rather that the high relative abundance of *P. agricolaris* corresponds with a reduction in microbiota phylogenetic diversity (Figure 4a).

**Figure 5.**
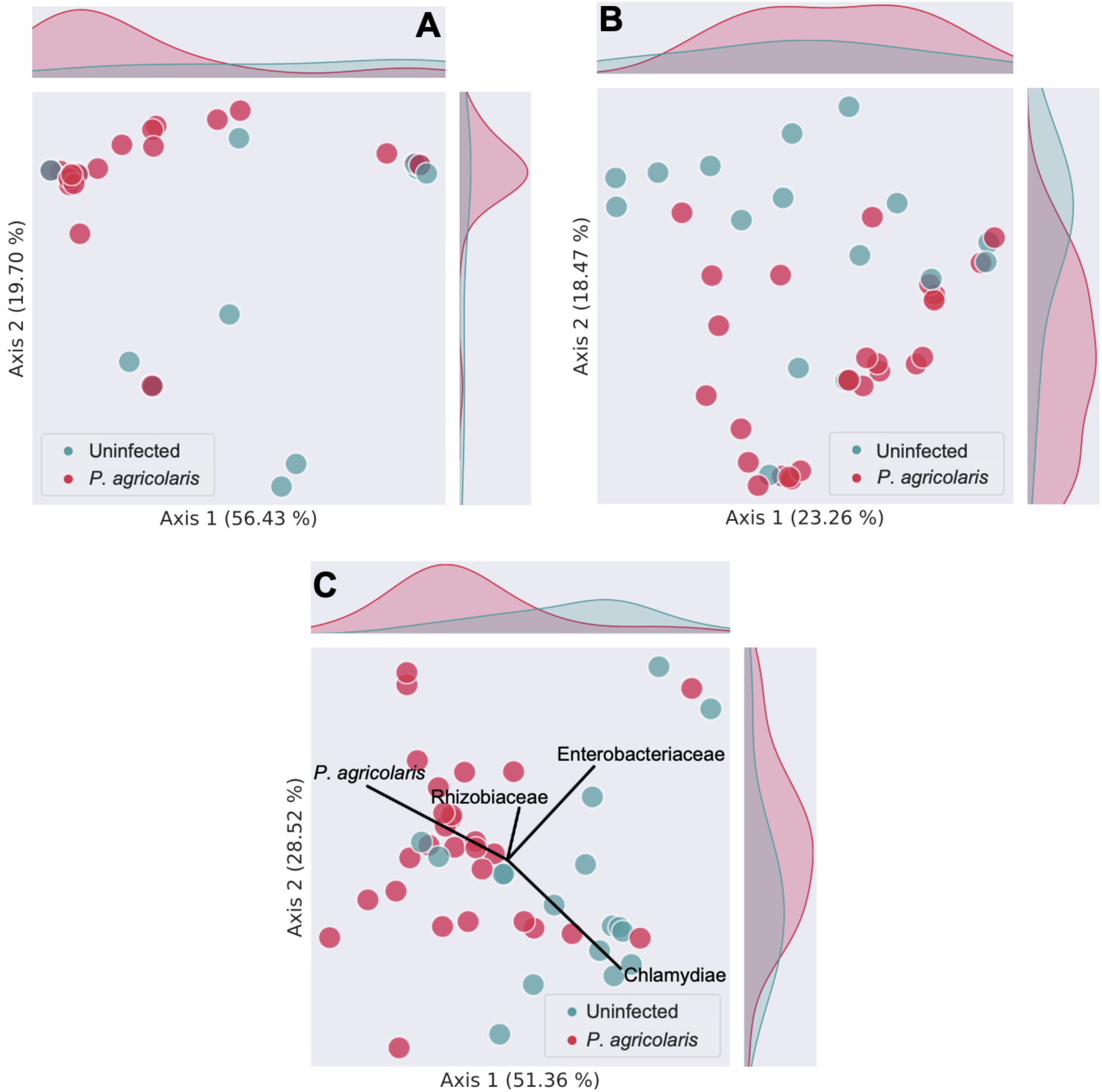
High degrees of *P. agricolaris* abundance results in similar microbiota compositions among infected amoebae. Beta-diversity ordination plots comparing the microbiota of *P. agricolaris* infected and uninfected *D. discoideum*, where each point represents an individual isolates microbiota and the closer points are to each other, the more similar their microbial communities. Kernel density estimate plots along the x and y axes indicate the distribution of points along the respective axes and highlight each groups concentration in ordination space. A) Weighted UniFrac PCoA plot showing the difference between infected and uninfected microbiota structures. B) Unweighted UniFrac PCoA showing the difference in microbiota compositions. C) Aitchison PCA plot showing the contribution of key bacterial taxa to sample clustering.

The contribution of *P. agricolaris* to the similarity between infected *D. discoideum* samples is more quantitatively highlighted via Robust Aitchison analysis, which is a novel beta-diversity metric that is more adept at handling the sparsity of microbial community data. Here, Aitchison analysis uses centered log ratio transformations as opposed to relative abundances (which can often yield spurious results). Pairing log ratio transformation with the compositional Aitchison distances allows for better interpretations of microbiome composition and also yields insight into salient members of microbial communities that are responsible for sample clustering [60]. Along with showing significant differences between *P. agricolaris* infected and uninfected *D. discoideum* microbiota (PERMANOVA, pseudo-F=9.28284, p=0.001; PERMDISP, F=0.655692, p=0.392) Robust Aitcheson PCA highlights the influence of *P. agricolaris* (indicated by labeled vector) on the grouping of infected *D. discoideum* microbiota (Figure 5c).

*D. discoideum* inoculated with *Paraburkholderia* in the laboratory showed similar, yet more pronounced, differences when compared to uninfected *D. discoideum. D. discoideum* microbiota structure, determined using weighted UniFrac, was significantly altered by infection with *P. agricolaris, P. hayleyella*, or *P. bonniea* when compared to uninfected *D. discoideum* (PERMANOVA, pseudo-F ≥ 7.384689, p ≤ 0.007, PERMDISP, F=26.8176, p=0.001 where significant PERMDISP results were negated by clear PCoA grouping) (Figure 6a). Likewise, *D. discoideum* microbiota composition, evaluated using unweighted UniFrac, was altered by infection with either *P. agricolaris, P. hayleyella*, or *P. bonniea* (PERMANOVA, pseudo-F ≥ 4.200470, p ≤ 0.002, PERMDISP, F=2.30245, p=0.11) (Figure 6b). Microbiota compositional comparisons using Aitchison distances showed a clear distinction between the microbiota of *Paraburkholderia* infected amoebae and uninfected amoebae (PERMANOVA, pseudo-F=22.282, p=0.001, PERMDISP, F=8.9081, p=0.001). Furthermore, Aitchison PCA highlights the contribution of each *Paraburkholderia* species to microbiota composition (Figure 6c). These results are consistent with the reduction in evenness, richness, and phylogenetic diversity observed in α-diversity comparisons.

**Figure 6.**
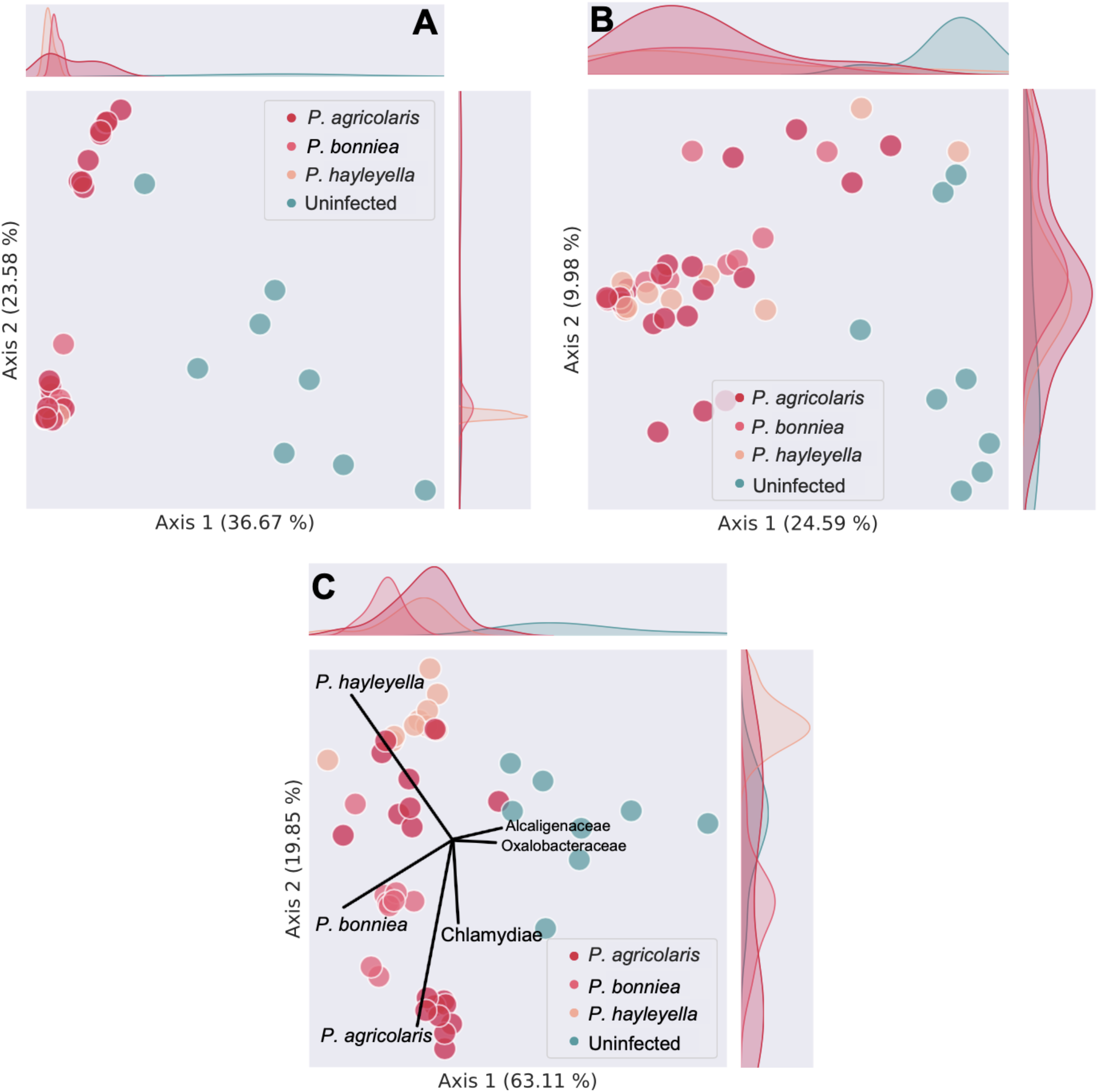
Over-abundance of each *Paraburkholderia* species in the microbiota results in similar microbiota compositions among infected amoebae. Beta-diversity ordination plots comparing the microbiota of *D. discoideum* artificially infected with each *Paraburkholderia* species to uninfected *D. discoideum*, where each point represents an individual isolates microbiota and the closer points are to each other, the more similar their microbial communities. Kernel density estimate plots along the x and y axes indicate the distribution of points along the respective axes and highlight each groups concentration in ordination space. A) Weighted UniFrac PCoA plot showing the difference between infected and uninfected microbiota structures. B) Unweighted UniFrac PCoA showing the difference in microbiota compositions. C) Aitchison PCA plot showing the contribution of key bacterial taxa to sample clustering.

### *P. agricolaris* positively associates with *Rhizobiales* in the natural *D. discoideum* microbiota

Although we did not find indication of an enhancement in bacterial food carriage by *Paraburkholderia* via compositional comparisons, we reasoned that food carriage could be indicated by specific co-associations with *P. agricolaris*. Therefore, we performed co-association analyses using the SparCC method, which accounts for bacterial presence and abundance, and is adept at handling the sparsity of microbial community data. We identified only one positive association between *P. agricolaris*, represented by the order Burkholderiales, and the order Rhizobiales (p=0.0297), and no significant negative association. Overall, significant associations between other members of the amoeba microbiota were scarce (Figure 7).

**Figure 7.**
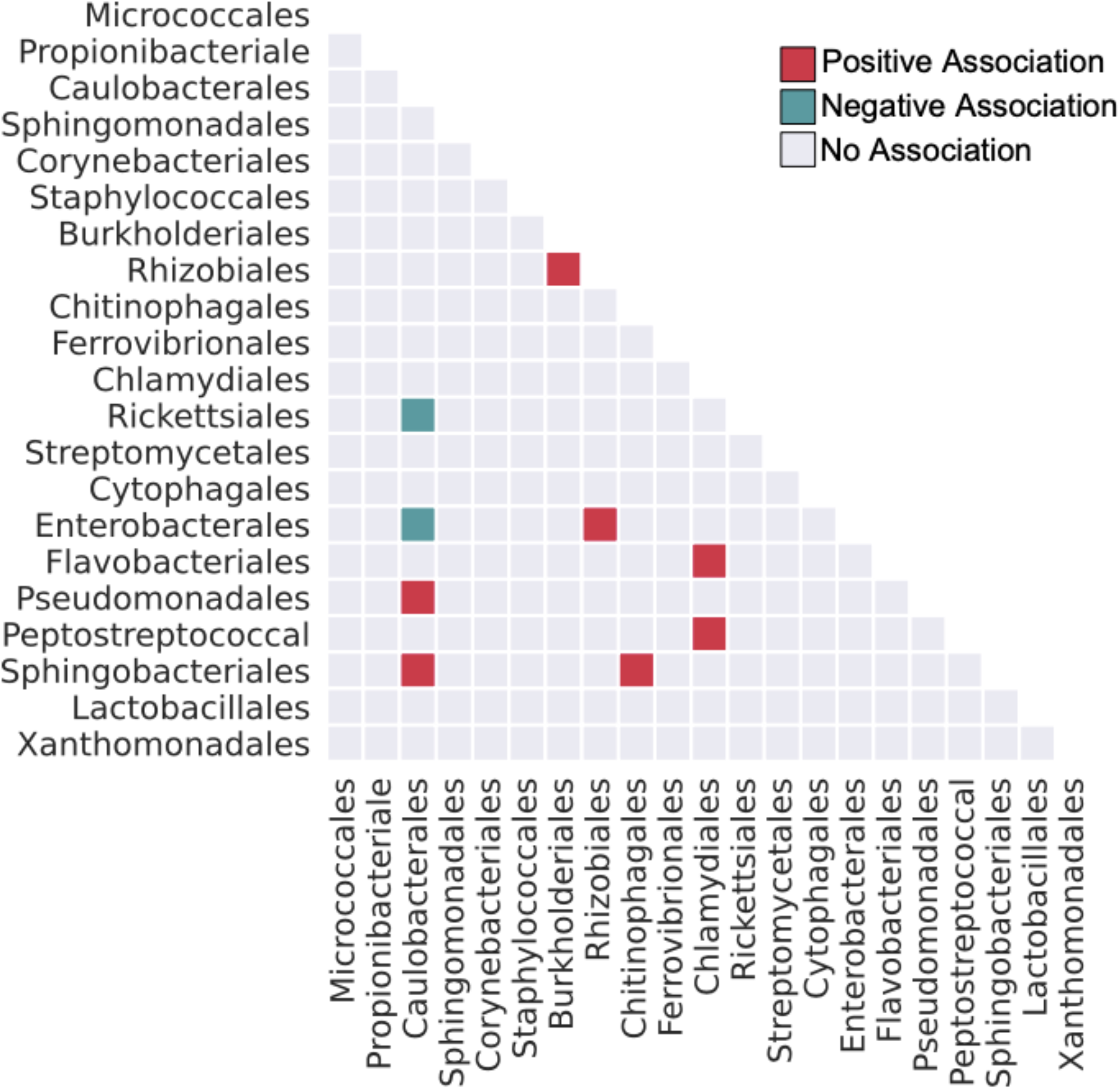
*P. agricolaris* positively associates with members of the order Rhizobiales. Heatmap showing significant co-associations between bacterial orders within the natural *D. discoideum* microbiota. *P. agricolaris* is represented by the order Burkholderiales.

### Chlamydiae is abundant in the natural *D. discoideum* microbiota but does not affect its diversity

Given the high prevalence of Chlamydiae in *D. discoideum* populations, we investigated how it might impact the natural *D. discoideum* microbiota. We found that the phylum Chlamydiae accounted for 21.57% of all sequence reads, making it the second most abundant group. The average abundance of Chlamydiae across infected samples was 43.34%, where Chlamydiae accounted for 36.86% of all reads in infected samples and for greater than 90% of reads in 8/25 infected samples (Figure 3a). Chlamydiae infection was not correlated with microbiota evenness, richness, or phylogenetic diversity (Pielou’s Evenness, p=0.731175; observed features (ASVs), p=0.257608; Faith’s Phylogenetic Diversity, p=0.213492). However, Chlamydiae infected *D. discoideum* did have distinct microbiota compositions and structures, measured by Weighted UniFrac (PERMANOVA, pseudo-F=5.38621, p=0.004, PERMDISP, F=0.164404, p=0.660) and unweighted UniFrac (PERMANOVA, pseudo-F=4.31874, p=0.003, PERMDISP, F=8.00349, p=0.011) respectively.

## Discussion

While previous studies of the *Paraburkholderia – D. discoideum* symbiosis have worked with samples isolated from nature, exploration of the farming phenomenon has been primarily laboratory based. Therefore, we aimed to investigate this interaction as it might occur in a more natural context. It has been previously established that *Paraburkholderia* is prevalent in *D. discoideum* populations (Haselkorn et al., 2019). Here, we expand upon these findings by quantifying *Paraburkholderia* prevalence nearly twenty years after an initial sampling. Our recent survey of natural *D. discoideum* populations showed little to no incidences of *P. bonniea* and *P. hayleyella* infection, yet *P. agricolaris* remained prevalent. Furthermore, we found evidence that *P. agricolaris* is environmentally acquired in nature, which is supported by its high degree of genetic diversity. While the presumed benefit of farming could drive this symbionts’ persistence, overall, we did not find support for our hypothesis that the *Paraburkholderia* farming symbiont would increase the diversity of *D. discoideum* microbiota. Rather, *P. agricolaris* tended to infectiously dominate the *D. discoideum* microbiota in nature. Artificial inoculation of each farming symbiont recapitulated this pattern, where *P. bonniea* and *P. hayleyella* also infectiously dominated the *D. discoideum* microbiota, making it unclear if food carriage is ecologically relevant.

It is a challenge to quantify the farming phenotype in nature, as measurement of the phenotype has primarily been done in the lab. Laboratory tests of farming are generally in a controlled environment, starting with *D. discoideum* clones being passaged and cleared of natural transient bacteria and then provided with an ample supply of specific types of food bacteria (Brock et al., 2011). *Paraburkholderia* confers secondary food carriage under these conditions, increasing the microbial diversity. This food carriage is mostly extracellular and varies by *Paraburkholderia* genotype and food bacterial species (Khojandi et al., 2019; Miller et al., 2020). Recent work has shown that *Paraburkholderia* symbionts inhibit phagosome acidification (Tian et al., 2022), suggesting that food carriage could be mediated by the creation of a more hospitable environment for other bacteria that get ingested. In the soil environment, however, many fruiting bodies contain microbiota, both with and without *Paraburkholderia* (Brock et al., 2018; Sallinger et al., 2021), and in our study, *Paraburkholderia* did not increase the diversity of this microbiota. In fact, in our lab infection assay, where fruiting bodies were collected directly from the microbially diverse soil environment, *Paraburkholderia* significantly decreased overall diversity.

Other effects that *Paraburkholderia* have on the microbiota are less clear. Although *Paraburkholderia* does not increase the diversity of the microbiome, we cannot rule out the possibility that it causes an increase in overall microbial load, which could allow an increase in other edible bacteria. We did observe that *Paraburkholderia* infected amoebae had compositionally similar microbiota, which could be consistent with farming if *Paraburkholderia* favored the carriage of particular bacteria more likely to be food. These compositional changes, however, were most likely due to a decrease in phylogenetic diversity correlated with *Paraburkholderia* infection. Given the abundant natural microbiota of *D. discoideum* and other social amoeba species in the absence of *Paraburkholderia* infection, the potential benefits of food carriage are likely not specific to *Paraburkholderia* infected amoebas. The only other significant effect of *Paraburkholderia* on the microbiota was its positive co-association with Rhizobiales. While the implications of this association are unclear, it merits further investigation, given observations of similar patterns in experimental studies (Khojandi et al., 2019) and the importance of Rhizobiales in the soil environment.

*Paraburkholderia* may have alternative benefits which increase host fitness. A previous study has demonstrated that *D. discoideum* spore production is significantly reduced when exposed to the toxin ethidium bromide but is not impacted under the same conditions when harboring *Paraburkholderia* (Brock et al., 2016). While this study suggested that farming associated bacteria over-compensated for fewer sentinel cells produced by *Paraburkholderia* infected hosts, the dominance of *Paraburkholderia* in the amoeba microbiota could suggest *Paraburkholderia* itself plays a role in detoxification. Such roles are often assumed by *Burkholderia* symbionts of both insects and fungi. For example, the bean bug *Riptortus pedestris* harbors *Burkholderia* symbionts that protect their host from the harmful effects of the insecticide fenitrothion, and other *Burkholderia* symbionts protect their fungal hosts from a variety of anti-fungal agents (Kikuchi et al., 2012; Nazir et al., 2014). Alternatively, it is possible that *P. agricolaris* competitively excludes other potentially detrimental bacteria from infecting the amoeba fruiting body, just as the certain *Burkholderia* symbionts out compete non-native symbionts in the crypt of the bean bug *Riptortus pedestris* (Itoh et al., 2019).

There is conflicting evidence as to whether these *Paraburkholderia-D. discoideum* symbioses are co-evolved mutualisms. The prevalence of *P. agricolaris* in nature is consistent with beneficial effects that outweighs any cost of infection, as would be seen in a mutualism. However, this high degree of prevalence could also be attributed to a more parasitic relationship. Likewise, finding almost no incidence of *P. bonniea* or *P. hayleyella* infection could indicate their detriment or lack of relevance to host fitness. More time points would be needed to establish this trend ecologically. Indeed, some *P. hayleyella* have a demonstrably large fitness cost in laboratory assays, though *P. bonniea* has not been reported to significantly impact *D. discoideum* spore production (Haselkorn et al., 2019; Khojandi et al., 2019). Many long term co-evolved mutualisms lack horizontal transmission, and the symbiont genomes degrade as they adapt to living solely inside their host and vertical transmission (Moran, 1996). However, *P. agricolaris* possesses a larger genome than both *P. bonniea* and *P. hayleyella* (Brock et al., 2020). Due to this larger genome, *P. agricolaris* can utilize a wider diversity of carbon sources than its counterparts, which makes it more similar to other free-living *Paraburkholderia* and *Burkholderia* relatives. It also infects a fewer proportion of spores than the other *Paraburkholderia* symbiont species (Khojandi et al., 2019; Miller et al., 2020), suggesting that it is not strongly dependent on vertically transmission. *P. bonniea* and *P. hayleyella*, which infect a higher proportion of spores in laboratory experiments, may be more reliant on vertical transmission in nature as their smaller genomes suggest they may not be as capable of living freely in the soil.

It is also possible that *P. agricolaris* is simply prevalent in *D. discoideum* populations because it is prevalent in, and subsequently acquired from, the soil environment. This is supported by the findings of Sallinger *et al*. (Sallinger et al., 2021), where *P. agricolaris* was detectable in relatively high frequency in bulk soil sample from which amoebae were cultured. Environmental acquisition could explain the high genetic diversity of *P. agricolaris* in natural populations, which differs from the pattern of a single beneficial haplotype sweeping through a population after initial infection, as often occurs in insect symbionts (Cockburn et al., 2013; Himler et al., 2011). Here, *P. agricolaris* would not be subjected to the population bottleneck and subsequent loss of genetic diversity elicited by vertical transmission. Additionally, a wider diversity of *P. agricolaris* haplotypes could be acquired from the soil upon initial infection. The intracellular and extracellular occupation of *P. agricolaris* makes it both horizontally and vertically transmissible (Khojandi et al., 2019), a pattern that has been observed in insects, where the oriental chinch bug obtains its *Burkholderia* symbiont from both parent and environment (Itoh et al., 2014). It is also consistent with the observation of these same *Paraburkholderia* haplotypes infecting other, distantly related co-occurring species of social amoebae. This acquisition could be driven by either the host (Rashidi & Ostrowski, 2019) or bacteria (Shu et al., 2018). The ease of environmental acquisition of *P. agricolaris* may ultimately lessen the pressure to develop and maintain vertical transmission and limit co-evolution. Such patterns have also been observed in symbioses between *Burkholderia* and broad-headed insects, where acquisition of *Burkholderia* from the environment dampens co-evolutionary dynamics (Garcia et al., 2014; Kikuchi et al., 2007).

Recent evidence has also suggested that environmental factors could impact *Paraburkholderia* prevalence, where *P. hayleyella* infection provided additional detriment to *D. discoideum* under artificial warming conditions (Shu et al., 2020). Therefore, our findings regarding symbiont prevalence may not be generalizable to other *D. discoideum* populations occurring in different soil environments. However, the environmental parameters that impact this symbiosis merit further investigation. Even within our sub-sampling the symbiont prevalence was highly variable, suggesting that differences in microhabitat may have large effects. Additional sampling of different geographic locations at different times of year may help to elucidate these patterns.

Like *Paraburkholderia*, we found that Chlamydiae is highly prevalent in natural populations, though the reasons for this are less clear. Laboratory studies of Chlamydiae in *D. discoideum* found no fitness cost of infection (Haselkorn et al., 2021), but potential benefits have yet to be explored. Like *Paraburkholderia*, Chlamydiae was overabundant in the microbiota that it occupied, resulting in compositional uniformity. Co-association analyses within the microbiota showed positive associations between the order Chlamydiales and the orders Flavobacteriales and Peptostreptococcal, suggesting that this bacterium could be altering the social amoeba microbiota in meaningful ways. The high prevalence of Chlamydiae in natural populations suggests an ecological relevance for this symbiont that is worth further studying.

Overall, the variability regarding *P. agricolaris* localization, prevalence, transmission patterns, genetic diversity, and costs of infection might indicate movement along the parasitism – mutualism spectrum. Such dynamics can be difficult to study in more complex systems. Thus, the *D. discoideum* – *Paraburkholderia* symbiosis, as well as the *D. discoideum* microbiota as a whole, offers a useful model for understanding symbiotic development. Future studies that aim to identify specific environmental parameters and soil conditions that either facilitate or hinder this association will be useful in providing further ecological contextualization. Furthermore, future studies that aim to investigate additional functions of *Paraburkholderia* will further our understanding of the context dependent nature of symbiotic interactions in general.

## Acknowledgments

We would like to thank Julia Roberson for technical assistance with the laboratory inoculation experiment, Susanne DiSalvo for valuable discussions and comments on the manuscript, and Joan Strassmann for her feedback on this work. Funding was provided by the University of Central Arkansas (UCA) Advancement of Undergraduates in Research in the Science (AURS) program, the Arkansas Department of Higher Education Student Undergraduate Research Fellowship (SURF), and the UCA College of Natural Science and Mathematics. Bioinformatics were performed on the UCA Dell Blade Server System, “BRUCE,” which was funded by NSF Award Number 1726009. This project has been supported by the Arkansas INBRE program, with a grant from the National Institute of General Medical Sciences (NIGMS), P20 GM103429 from the National Institute of Health.

## Data Accessibility

All sequences used to confirm *Paraburkholderia* infection and identity in natural *D. discoideum* populations have been uploaded to GenBank under accession numbers MZ395652 - MZ395810. Sequences used for *P. agricolaris* haplotype analyses have been uploaded under accession numbers MZ399634 - MZ399696. Next-generation 16S rRNA sequences obtained for microbiota comparisons have been uploaded as BioProject PRJNA737516. Sequences used to confirm amoeba species identity were uploaded to GenBank under accession numbers MZ396209-MZ396243.

## Author Contributions

TSH and JGD designed research. TSH, MH, HO, and JGD performed research. JGD and MSR analyzed data. TSH and JGD wrote the paper with edits from MH and MSR.

## Supplemental Tables

**Supplemental Table 1:**
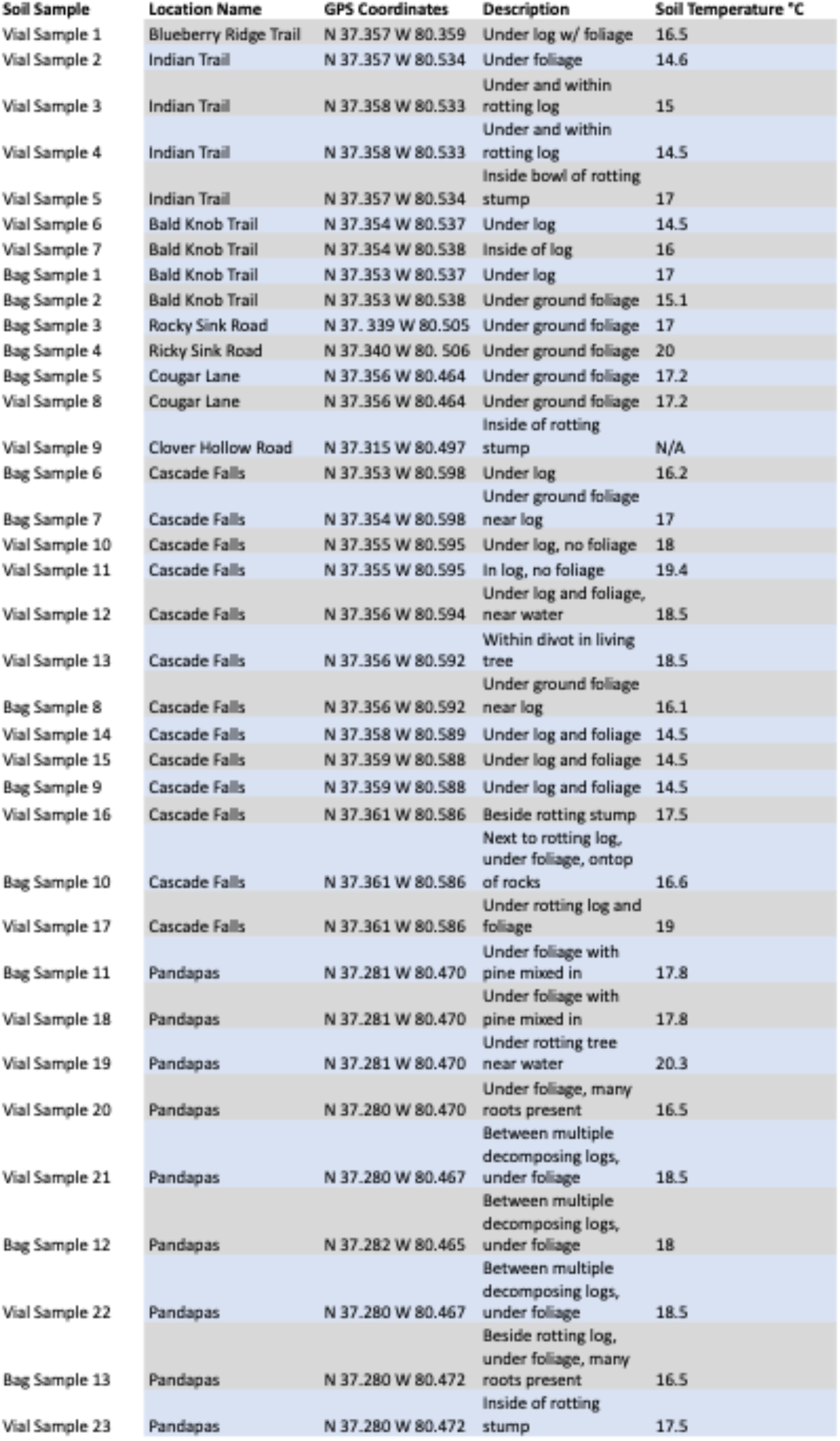
Soil Sample Metadata

**Supplemental Table 2:**
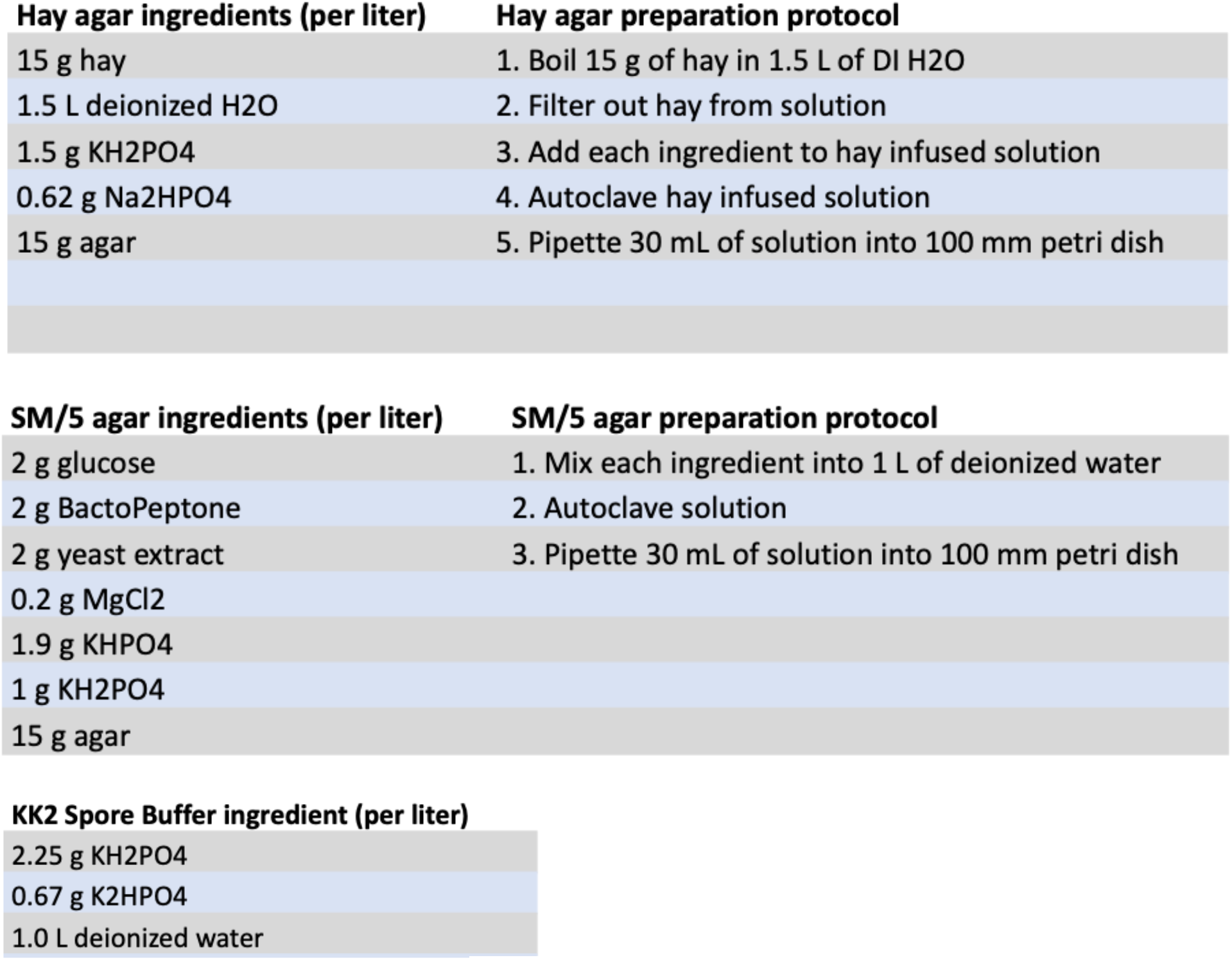
Microbiological Media Preparation and KK2 extraction buffer

**Supplemental Table 3:**
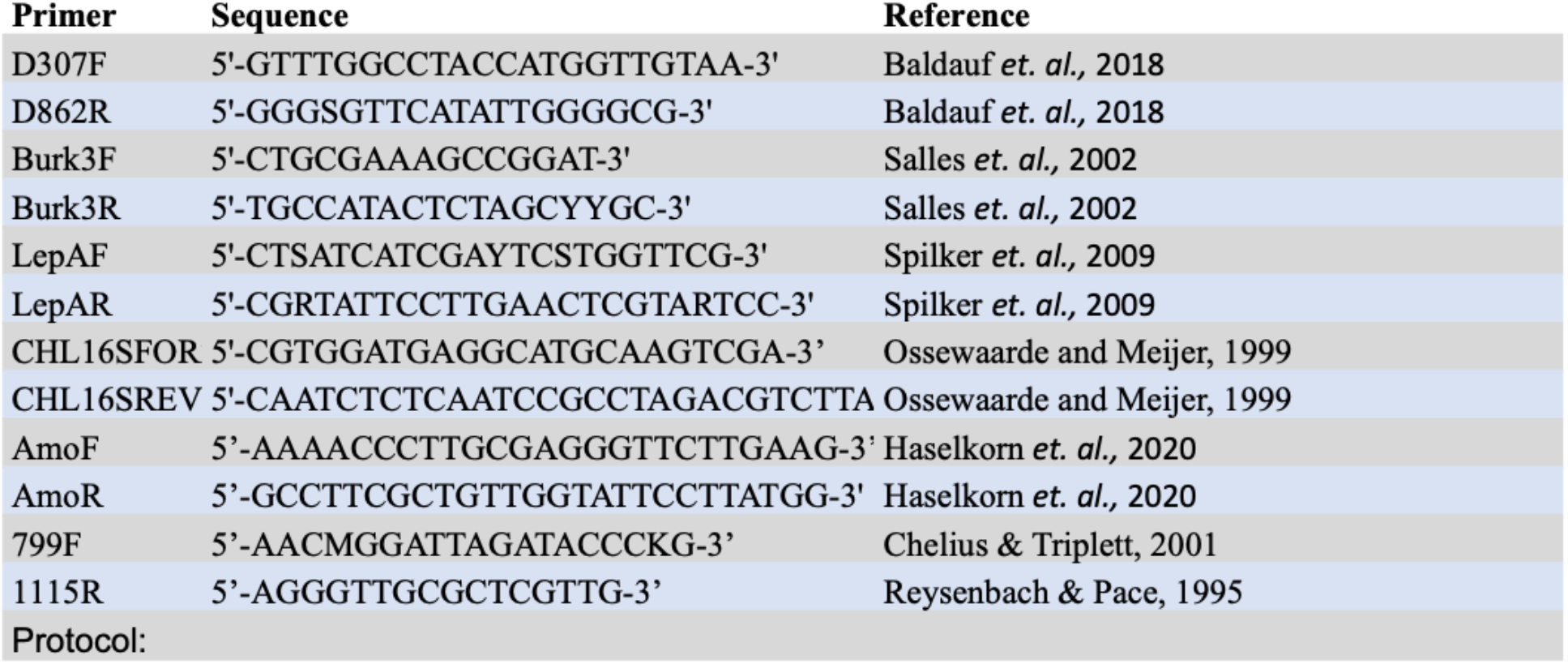
Primer Sets

**Supplemental Table 4:**
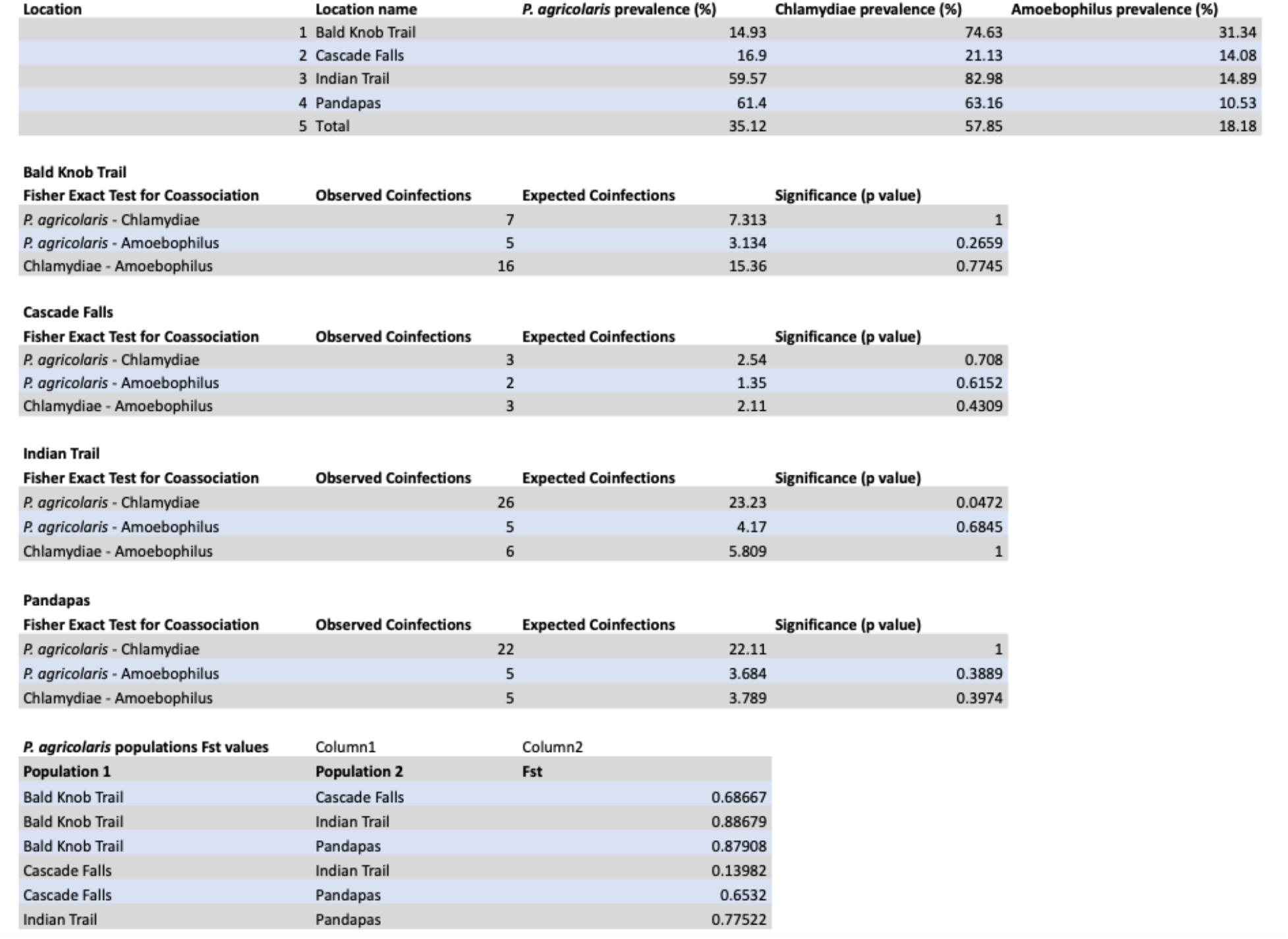
Prevalence, co-associations, and genetic community comparisons of symbiont screenings

| Location | Location name | *P. agricularis* prevalence (%) | Chlamydiae prevalence (%) | Amoebophilus prevalence (%) |
| --- | --- | --- | --- | --- |
|  | 1 Bald Knob Trail | 14.93 | 74.63 | 31.34 |
|  | 2 Cascade Falls | 16.9 | 21.13 | 14.08 |
|  | 3 Indian Trail | 59.57 | 82.98 | 14.89 |
|  | 4 Pandapas | 61.4 | 63.16 | 10.53 |
|  | 5 Total | 35.12 | 57.85 | 18.18 |

**Bald Knob Trail**
| Fisher Exact Test for Coassociation | Observed Coinfections | Expected Coinfections | Significance (p value) |
| --- | --- | --- | --- |
| *P. agricularis* - Chlamydiae | 7 | 7.313 | 1 |
| *P. agricularis* - Amoebophilus | 5 | 3.134 | 0.2659 |
| Chlamydiae - Amoebophilus | 16 | 15.36 | 0.7745 |

**Cascade Falls**
| Fisher Exact Test for Coassociation | Observed Coinfections | Expected Coinfections | Significance (p value) |
| --- | --- | --- | --- |
| *P. agricularis* - Chlamydiae | 3 | 2.54 | 0.708 |
| *P. agricularis* - Amoebophilus | 2 | 1.35 | 0.6152 |
| Chlamydiae - Amoebophilus | 3 | 2.11 | 0.4309 |

**Indian Trail**
| Fisher Exact Test for Coassociation | Observed Coinfections | Expected Coinfections | Significance (p value) |
| --- | --- | --- | --- |
| *P. agricularis* - Chlamydiae | 26 | 23.23 | 0.0472 |
| *P. agricularis* - Amoebophilus | 5 | 4.17 | 0.6845 |
| Chlamydiae - Amoebophilus | 6 | 5.809 | 1 |

**Pandapas**
| Fisher Exact Test for Coassociation | Observed Coinfections | Expected Coinfections | Significance (p value) |
| --- | --- | --- | --- |
| *P. agricularis* - Chlamydiae | 22 | 22.11 | 1 |
| *P. agricularis* - Amoebophilus | 5 | 3.684 | 0.3889 |
| Chlamydiae - Amoebophilus | 5 | 3.789 | 0.3974 |

| *P. agricularis* populations Fst values | Column1 | Column2 |
| --- | --- | --- |
| Population 1 | Population 2 | Fst |
| Bald Knob Trail | Cascade Falls | 0.68667 |
| Bald Knob Trail | Indian Trail | 0.88679 |
| Bald Knob Trail | Pandapas | 0.87908 |
| Cascade Falls | Indian Trail | 0.13982 |
| Cascade Falls | Pandapas | 0.6532 |
| Indian Trail | Pandapas | 0.77522 |

**Supplemental Table 5:** Soil samples and corresponding *P. agricolaris* haplotypes

| Location | Soil Sample Code | *P. agriculturalis* haplotypes present |
| --- | --- | --- |
| Bald Knob Trail | 1B | 10 |
| Cascade Falls | 2B1 | 5,6,7 |
| Indian Trail | 3A1 | 5,6,7 |
| Indian Trail | 3A2 | 7 |
| Indian Trail | 3B | 7,8 |
| Pandapas | 4A1 | 1,2,3,4,5 |
| Pandapas | 4A2 | 1,3,4 |
| Pandapas | 4A3 | 4 |

## References

Anderson, M. J. (2001). A new method for non-parametric multivariate analysis of variance in ecology. https://doi.org/10.1111/J.1442-9993.2001.01070.PP.X

Anderson, R. M., & May, R. M. (1982). Coevolution of hosts and parasites. Parasitology, 85 (Pt 2), 411–426. https://doi.org/10.1017/s0031182000055360

Baldauf, S. L., Romeralo, M., Fiz-Palacios, O., & Heidari, N. (2018). A Deep Hidden Diversity of Dictyostelia. Protist, 169(1), 64–78. https://doi.org/10.1016/j.protis.2017.12.005

Bokulich, N. A., Kaehler, B. D., Rideout, J. R., Dillon, M., Bolyen, E., Knight, R., Huttley, G. A., & Gregory Caporaso, J. (2018). Optimizing taxonomic classification of marker-gene amplicon sequences with QIIME 2’s q2-feature-classifier plugin. Microbiome, 6(1), 90. https://doi.org/10.1186/s40168-018-0470-z

Bozzaro, S. (2019). The past, present and future of Dictyostelium as a model system. The International Journal of Developmental Biology, 63(8-9–10), 321–331. https://doi.org/10.1387/ijdb.190128sb

Brock, D. A., Callison, W.É., Strassmann, J. E., & Queller, D. C. (2016). Sentinel cells, symbiotic bacteria and toxin resistance in the social amoeba Dictyostelium discoideum. Proceedings. Biological Sciences, 283(1829), 20152727. https://doi.org/10.1098/rspb.2015.2727

Brock, D. A., Douglas, T. E., Queller, D. C., & Strassmann, J. E. (2011). Primitive agriculture in a social amoeba. Nature, 469(7330), 393–396. https://doi.org/10.1038/nature09668

Brock, D. A., Haselkorn, T. S., Garcia, J. R., Bashir, U., Douglas, T. E., Galloway, J., Brodie, F., Queller, D. C., & Strassmann, J. E. (2018). Diversity of Free-Living Environmental Bacteria and Their Interactions With a Bactivorous Amoeba. Frontiers in Cellular and Infection Microbiology, 8, 411. https://doi.org/10.3389/fcimb.2018.00411

Brock, D. A., Noh, S., Hubert, A. N. M., Haselkorn, T. S., DiSalvo, S., Suess, M. K., Bradley, A. S., Tavakoli-Nezhad, M., Geist, K. S., Queller, D. C., & Strassmann, J. E. (2020). Endosymbiotic adaptations in three new bacterial species associated with Dictyostelium discoideum: Paraburkholderia agricolaris sp. nov., Paraburkholderia hayleyella sp. nov., and Paraburkholderia bonniea sp. nov. PeerJ, 8, e9151. https://doi.org/10.7717/peerj.9151

Brüssow, H. (2007). Bacteria between protists and phages: From antipredation strategies to the evolution of pathogenicity. Molecular Microbiology, 65(3), 583–589. https://doi.org/10.1111/j.1365-2958.2007.05826.x

Callahan, B. J., McMurdie, P. J., Rosen, M. J., Han, A. W., Johnson, A. J. A., & Holmes, S. P. (2016). DADA2: High-resolution sample inference from Illumina amplicon data. Nature Methods, 13(7), 581–583. https://doi.org/10.1038/nmeth.3869

Camacho, C., Coulouris, G., Avagyan, V., Ma, N., Papadopoulos, J., Bealer, K., & Madden, T. L. (2009). BLAST+: Architecture and applications. BMC Bioinformatics, 10, 421. https://doi.org/10.1186/1471-2105-10-421

Casadevall, A. (2008). Evolution of intracellular pathogens. Annu Rev Microbiol, 62, 19–33. https://doi.org/10.1146/annurev.micro.61.080706.093305

Chang, Q., Luan, Y., & Sun, F. (2011). Variance adjusted weighted UniFrac: A powerful beta diversity measure for comparing communities based on phylogeny. BMC Bioinformatics, 12, 118. https://doi.org/10.1186/1471-2105-12-118

Chen, G., Zhuchenko, O., & Kuspa, A. (2007). Immune-like phagocyte activity in the social amoeba. Science, 317(5838), 678–681. https://doi.org/10.1126/science.1143991

Chen, J., Bittinger, K., Charlson, E. S., Hoffmann, C., Lewis, J., Wu, G. D., Collman, R. G., Bushman, F. D., & Li, H. (2012). Associating microbiome composition with environmental covariates using generalized UniFrac distances. Bioinformatics (Oxford, England), 28(16), 2106–2113. https://doi.org/10.1093/bioinformatics/bts342

Cockburn, S. N., Haselkorn, T. S., Hamilton, P. T., Landzberg, E., Jaenike, J., & Perlman, S. J. (2013). Dynamics of the continent-wide spread of a Drosophila defensive symbiont. Ecology Letters, 16(5), 609–616. https://doi.org/10.1111/ele.12087

Delafont, V., Samba-Louaka, A., Bouchon, D., Moulin, L., & Héchard, Y. (2015). Shedding light on microbial dark matter: A TM6 bacterium as natural endosymbiont of a free-living amoeba. Environmental Microbiology Reports, 7(6), 970–978. https://doi.org/10.1111/1758-2229.12343

Dinh, C., Farinholt, T., Hirose, S., Zhuchenko, O., & Kuspa, A. (2018). Lectins modulate the microbiota of social amoebae. Science (New York, N.Y.), 361(6400), 402–406. https://doi.org/10.1126/science.aat2058

DiSalvo, S., Haselkorn, T. S., Bashir, U., Jimenez, D., Brock, D. A., Queller, D. C., & Strassmann, J. E. (2015). Burkholderia bacteria infectiously induce the proto-farming symbiosis of Dictyostelium amoebae and food bacteria. Proceedings of the National Academy of Sciences of the United States of America, 112(36), E5029–5037. https://doi.org/10.1073/pnas.1511878112

Dunn, J. D., Bosmani, C., Barisch, C., Raykov, L., Lefrançois, L. H., Cardenal-Muñoz, E., López-Jiménez, A. T., & Soldati, T. (2018). Eat Prey, Live: Dictyostelium discoideum As a Model for Cell-Autonomous Defenses. Frontiers in Immunology, 8. https://doi.org/10.3389/fimmu.2017.01906

Esmaeel, Q., Miotto, L., Rondeau, M., Leclère, V., Clément, C., Jacquard, C., Sanchez, L., & Barka, E. A. (2018). Paraburkholderia phytofirmans PsJN-Plants Interaction: From Perception to the Induced Mechanisms. Frontiers in Microbiology, 9, 2093. https://doi.org/10.3389/fmicb.2018.02093

Ewald, P. W. (1987). Transmission modes and evolution of the parasitism-mutualism continuum. Annals of the New York Academy of Sciences, 503, 295–306. https://doi.org/10.1111/j.1749-6632.1987.tb40616.x

Faith, D. P., Minchin, P. R., & Belbin, L. (1987). Compositional Dissimilarity as a Robust Measure of Ecological Distance. Vegetatio, 69(1/3), 57–68.

Farinholt, T., Dinh, C., & Kuspa, A. (2019). Microbiome management in the social amoeba Dictyostelium discoideum compared to humans. The International Journal of Developmental Biology, 63(8-9–10), 447–450. https://doi.org/10.1387/ijdb.190240ak

Garcia, J. R., Laughton, A. M., Malik, Z., Parker, B. J., Trincot, C. S L, Chiang, S., Chung, E., & Gerardo, N. M. (2014). Partner associations across sympatric broad-headed bug species and their environmentally acquired bacterial symbionts. Molecular Ecology, 23(6), 1333–1347. https://doi.org/10.1111/mec.12655

Greub, G., & Raoult, D. (2004). Microorganisms Resistant to Free-Living Amoebae. Clinical Microbiology Reviews, 17(2), 413–433. https://doi.org/10.1128/cmr.17.2.413-433.2004

Haselkorn, T. S., DiSalvo, S., Miller, J. W., Bashir, U., Brock, D. A., Queller, D. C., & Strassmann, J. E. (2019). The specificity of Burkholderia symbionts in the social amoeba farming symbiosis: Prevalence, species, genetic and phenotypic diversity. Molecular Ecology, 28(4), 847–862. https://doi.org/10.1111/mec.14982

Haselkorn, T. S., Jimenez, D., Bashir, U., Sallinger, E., Queller, D. C., Strassmann, J. E., & DiSalvo, S. (2021). Novel Chlamydiae and Amoebophilus endosymbionts are prevalent in wild isolates of the model social amoeba Dictyostelium discoideum. Environmental Microbiology Reports, 13(5), 126–142. https://doi.org/10.1111/1758-2229.12985

Himler, A. G., Adachi-Hagimori, T., Bergen, J. E., Kozuch, A., Kelly, S. E., Tabashnik, B. E., Chiel, E., Duckworth, V. E., Dennehy, T. J., Zchori-Fein, E., & Hunter, M. S. (2011). Rapid spread of a bacterial symbiont in an invasive whitefly is driven by fitness benefits and female bias. Science (New York, N.Y.), 332(6026), 254–256. https://doi.org/10.1126/science.1199410

Horn, M. (2008). Chlamydiae as symbionts in eukaryotes. Annual Review of Microbiology, 62, 113–131. https://doi.org/10.1146/annurev.micro.62.081307.162818

Horn, M., & Wagner, M. (2004). Bacterial endosymbionts of free-living amoebae. The Journal of Eukaryotic Microbiology, 51(5), 509–514. https://doi.org/10.1111/j.1550-7408.2004.tb00278.x

Ishida, K., Sekizuka, T., Hayashida, K., Matsuo, J., Takeuchi, F., Kuroda, M., Nakamura, S., Yamazaki, T., Yoshida, M., Takahashi, K., Nagai, H., Sugimoto, C., & Yamaguchi, H. (2014). Amoebal endosymbiont Neochlamydia genome sequence illuminates the bacterial role in the defense of the host amoebae against Legionella pneumophila. PloS One, 9(4), e95166. https://doi.org/10.1371/journal.pone.0095166

Itoh, H., Aita, M., Nagayama, A., Meng, X.-Y., Kamagata, Y., Navarro, R., Hori, T., Ohgiya, S., & Kikuchi, Y. (2014). Evidence of environmental and vertical transmission of Burkholderia symbionts in the oriental chinch bug, Cavelerius saccharivorus (Heteroptera: Blissidae). Applied and Environmental Microbiology, 80(19), 5974–5983. https://doi.org/10.1128/AEM.01087-14

Itoh, H., Jang, S., Takeshita, K., Ohbayashi, T., Ohnishi, N., Meng, X.-Y., Mitani, Y., & Kikuchi, Y. (2019). Host-symbiont specificity determined by microbe-microbe competition in an insect gut. Proceedings of the National Academy of Sciences of the United States of America, 116(45), 22673–22682. https://doi.org/10.1073/pnas.1912397116

Janssen, S., McDonald, D., Gonzalez, A., Navas-Molina, J. A., Jiang, L., Xu, Z. Z., Winker, K., Kado, D. M., Orwoll, E., Manary, M., Mirarab, S., & Knight, R. (2018). Phylogenetic Placement of Exact Amplicon Sequences Improves Associations with Clinical Information. MSystems, 3(3), e00021–18. https://doi.org/10.1128/mSystems.00021-18

Johnson, M., Zaretskaya, I., Raytselis, Y., Merezhuk, Y., McGinnis, S., & Madden, T. L. (2008). NCBI BLAST: A better web interface. Nucleic Acids Research, 36(Web Server issue), W5–9. https://doi.org/10.1093/nar/gkn201

Kaltenpoth, M., & Flórez, L. V. (2020). Versatile and Dynamic Symbioses Between Insects and Burkholderia Bacteria. Annual Review of Entomology, 65, 145–170. https://doi.org/10.1146/annurev-ento-011019-025025

Kearse, M., Moir, R., Wilson, A., Stones-Havas, S., Cheung, M., Sturrock, S., Buxton, S., Cooper, A., Markowitz, S., Duran, C., Thierer, T., Ashton, B., Meintjes, P., & Drummond, A. (2012). Geneious Basic: An integrated and extendable desktop software platform for the organization and analysis of sequence data. Bioinformatics (Oxford, England), 28(12), 1647–1649. https://doi.org/10.1093/bioinformatics/bts199

Kessin, R. H. (2001). Dictyostelium: Evolution, Cell Biology, and the Development of Multicellularity. Cambridge University Press.

Khojandi, N., Haselkorn, T. S., Eschbach, M. N., Naser, R. A., & DiSalvo, S. (2019). Intracellular Burkholderia Symbionts induce extracellular secondary infections; driving diverse host outcomes that vary by genotype and environment. The ISME Journal, 13(8), 2068–2081. https://doi.org/10.1038/s41396-019-0419-7

Kikuchi, Y., Hayatsu, M., Hosokawa, T., Nagayama, A., Tago, K., & Fukatsu, T. (2012). Symbiont-mediated insecticide resistance. Proceedings of the National Academy of Sciences of the United States of America, 109(22), 8618–8622. https://doi.org/10.1073/pnas.1200231109

Kikuchi, Y., Hosokawa, T., & Fukatsu, T. (2007). Insect-microbe mutualism without vertical transmission: A stinkbug acquires a beneficial gut symbiont from the environment every generation. Applied and Environmental Microbiology, 73(13), 4308–4316. https://doi.org/10.1128/AEM.00067-07

König, L., Wentrup, C., Schulz, F., Wascher, F., Escola, S., Swanson, M. S., Buchrieser, C., & Horn, M. (2019). Symbiont-Mediated Defense against Legionella pneumophila in Amoebae. MBio, 10(3), e00333–19. https://doi.org/10.1128/mBio.00333-19

Kruskal, W. H., & Wallis, W. A. (1952). Use of Ranks in One-Criterion Variance Analysis. Journal of the American Statistical Association, 47(260), 583–621. https://doi.org/10.2307/2280779

Kumar, S., Stecher, G., & Tamura, K. (2016). MEGA7: Molecular Evolutionary Genetics Analysis Version 7.0 for Bigger Datasets. Molecular Biology and Evolution, 33(7), 1870–1874. https://doi.org/10.1093/molbev/msw054

Lozupone, C. A., Hamady, M., Kelley, S. T., & Knight, R. (2007). Quantitative and qualitative beta diversity measures lead to different insights into factors that structure microbial communities. Applied and Environmental Microbiology, 73(5), 1576–1585. https://doi.org/10.1128/AEM.01996-06

Lozupone, C., & Knight, R. (2005). UniFrac: A new phylogenetic method for comparing microbial communities. Applied and Environmental Microbiology, 71(12), 8228–8235. https://doi.org/10.1128/AEM.71.12.8228-8235.2005

Maita, C., Matsushita, M., Miyoshi, M., Okubo, T., Nakamura, S., Matsuo, J., Takemura, M., Miyake, M., Nagai, H., & Yamaguchi, H. (2018). Amoebal endosymbiont Neochlamydia protects host amoebae against Legionella pneumophila infection by preventing Legionella entry. Microbes and Infection, 20(4), 236–244. https://doi.org/10.1016/j.micinf.2017.12.012

Martin, D. P., Murrell, B., Golden, M., Khoosal, A., & Muhire, B. (2015). RDP4: Detection and analysis of recombination patterns in virus genomes. Virus Evolution, 1(1), vev003. https://doi.org/10.1093/ve/vev003

Martino, C., Morton, J. T., Marotz, C. A., Thompson, L. R., Tripathi, A., Knight, R., & Zengler, K. (2019). A Novel Sparse Compositional Technique Reveals Microbial Perturbations. MSystems, 4(1), e00016–19. https://doi.org/10.1128/mSystems.00016-19

McDonald, D., Clemente, J. C., Kuczynski, J., Rideout, J. R., Stombaugh, J., Wendel, D., Wilke, A., Huse, S., Hufnagle, J., Meyer, F., Knight, R., & Caporaso, J. G. (2012). The Biological Observation Matrix (BIOM) format or: How I learned to stop worrying and love the ome-ome. GigaScience, 1(1), 7. https://doi.org/10.1186/2047-217X-1-7

McKinney, W. (2010). Data Structures for Statistical Computing in Python. https://doi.org/10.25080/MAJORA-92BF1922-00A

Miller, J. W., Bocke, C. R., Tresslar, A. R., Schniepp, E. M., & DiSalvo, S. (2020). Paraburkholderia Symbionts Display Variable Infection Patterns That Are Not Predictive of Amoeba Host Outcomes. Genes, 11(6), E674. https://doi.org/10.3390/genes11060674

Molmeret, M., Horn, M., Wagner, M., Santic, M., & Abu Kwaik, Y. (2005). Amoebae as training grounds for intracellular bacterial pathogens. Applied and Environmental Microbiology, 71(1), 20–28. https://doi.org/10.1128/AEM.71.1.20-28.2005

Moran, N. A. (1996). Accelerated evolution and Muller’s rachet in endosymbiotic bacteria. Proceedings of the National Academy of Sciences, USA, 93, 2873–2878.

Moran, N. A. (2006). Symbiosis. Curr Biol, 16(20), R866–71. https://doi.org/10.1016/j.cub.2006.09.019

Nazir, R., Tazetdinova, D. I., & van Elsas, J. D. (2014). Burkholderia terrae BS001 migrates proficiently with diverse fungal hosts through soil and provides protection from antifungal agents. Frontiers in Microbiology, 5, 598. https://doi.org/10.3389/fmicb.2014.00598

Okude, M., Matsuo, J., Nakamura, S., Kawaguchi, K., Hayashi, Y., Sakai, H., Yoshida, M., Takahashi, K., & Yamaguchi, H. (2012). Environmental chlamydiae alter the growth speed and motility of host acanthamoebae. Microbes and Environments, 27(4), 423–429. https://doi.org/10.1264/jsme2.me11353

Oliver, K. M., Campos, J., Moran, N. A., & Hunter, M. S. (2008). Population dynamics of defensive symbionts in aphids. Proceedings. Biological Sciences, 275(1632), 293–299. https://doi.org/10.1098/rspb.2007.1192

Ossewaarde, J. M., & Meijer, A. (1999). Molecular evidence for the existence of additional members of the order Chlamydiales. Microbiology (Reading, England), 145 (Pt 2), 411–417. https://doi.org/10.1099/13500872-145-2-411

Pielou, E. C. (1966). The measurement of diversity in different types of biological collections. https://doi.org/10.1016/0022-5193(66)90013-0

Raper, K. B. (2014). The Dictyostelids. Princeton University Press.

Rashidi, G., & Ostrowski, E. A. (2019). Phagocyte chase behaviours: Discrimination between Gram-negative and Gram-positive bacteria by amoebae. Biology Letters, 15(1), 20180607. https://doi.org/10.1098/rsbl.2018.0607

Redford, A. J., Bowers, R. M., Knight, R., Linhart, Y., & Fierer, N. (2010). The ecology of the phyllosphere: Geographic and phylogenetic variability in the distribution of bacteria on tree leaves. Environmental Microbiology, 12(11), 2885–2893. https://doi.org/10.1111/j.1462-2920.2010.02258.x

Robeson, M. S., O’Rourke, D. R., Kaehler, B. D., Ziemski, M., Dillon, M. R., Foster, J. T., & Bokulich, N. A. (2020). RESCRIPt: Reproducible sequence taxonomy reference database management for the masses. BioRxiv, 2020.10.05.326504. https://doi.org/10.1101/2020.10.05.326504

Rozas, J., Ferrer-Mata, A., Sánchez-DelBarrio, J. C., Guirao-Rico, S., Librado, P., Ramos-Onsins, S. E., & Sánchez-Gracia, A. (2017). DnaSP 6: DNA Sequence Polymorphism Analysis of Large Data Sets. Molecular Biology and Evolution, 34(12), 3299–3302. https://doi.org/10.1093/molbev/msx248

Salles, J. F., De Souza, F. A., & van Elsas, J. D. (2002). Molecular method to assess the diversity of Burkholderia species in environmental samples. Applied and Environmental Microbiology, 68(4), 1595–1603. https://doi.org/10.1128/AEM.68.4.1595-1603.2002

Sallinger, E., Robeson, M. S., & Haselkorn, T. S. (2021). Characterization of the bacterial microbiomes of social amoebae and exploration of the roles of host and environment on microbiome composition. Environmental Microbiology, 23(1), 126–142. https://doi.org/10.1111/1462-2920.15279

Sayers, E. W., Beck, J., Bolton, E. E., Bourexis, D., Brister, J. R., Canese, K., Comeau, D. C., Funk, K., Kim, S., Klimke, W., Marchler-Bauer, A., Landrum, M., Lathrop, S., Lu, Z., Madden, T. L., O’Leary, N., Phan, L., Rangwala, S. H., Schneider, V. A., … Sherry, S. T. (2021). Database resources of the National Center for Biotechnology Information. Nucleic Acids Research, 49(D1), D10–D17. https://doi.org/10.1093/nar/gkaa892

Schmitz-Esser, S., Toenshoff, E. R., Haider, S., Heinz, E., Hoenninger, V. M., Wagner, M., & Horn, M. (2008). Diversity of bacterial endosymbionts of environmental acanthamoeba isolates. Applied and Environmental Microbiology, 74(18), 5822–5831. https://doi.org/10.1128/AEM.01093-08

Shu, L., Qian, X., Brock, D. A., Geist, K. S., Queller, D. C., & Strassmann, J. E. (2020). Loss and resiliency of social amoeba symbiosis under simulated warming. Ecology and Evolution, 10(23), 13182–13189. https://doi.org/10.1002/ece3.6909

Shu, L., Zhang, B., Queller, D. C., & Strassmann, J. E. (2018). Burkholderia bacteria use chemotaxis to find social amoeba Dictyostelium discoideum hosts. The ISME Journal, 12(8), 1977–1993. https://doi.org/10.1038/s41396-018-0147-4

Spilker, T., Baldwin, A., Bumford, A., Dowson, C. G., Mahenthiralingam, E., & LiPuma, J. J. (2009). Expanded multilocus sequence typing for burkholderia species. Journal of Clinical Microbiology, 47(8), 2607–2610. https://doi.org/10.1128/JCM.00770-09

Sun, S., Noorian, P., & McDougald, D. (2018). Dual Role of Mechanisms Involved in Resistance to Predation by Protozoa and Virulence to Humans. Frontiers in Microbiology, 9, 1017. https://doi.org/10.3389/fmicb.2018.01017

Tian, Y., Peng, T., He, Z., Wang, L., Zhang, X., He, Z., & Shu, L. (2022). Symbiont-Induced Phagosome Changes Rather than Extracellular Discrimination Contribute to the Formation of Social Amoeba Farming Symbiosis. Microbiology Spectrum, e01727–21. https://doi.org/10.1128/spectrum.01727-21

Weiss, S., Xu, Z. Z., Peddada, S., Amir, A., Bittinger, K., Gonzalez, A., Lozupone, C., Zaneveld, J. R., Vázquez-Baeza, Y., Birmingham, A., Hyde, E. R., & Knight, R. (2017). Normalization and microbial differential abundance strategies depend upon data characteristics. Microbiome, 5(1), 27. https://doi.org/10.1186/s40168-017-0237-y

